# Clock proteins regulate spatiotemporal organization of clock genes to control circadian rhythms

**DOI:** 10.1101/2020.10.09.333732

**Authors:** Yangbo Xiao, Ye Yuan, Mariana Jimenez, Neeraj Soni, Swathi Yadlapalli

**Author notes:** These authors contributed equally to this paper (Yangbo Xiao, Ye Yuan).

## Abstract

Circadian clocks regulate ∼24 hour oscillations in gene expression, behavior, and physiology. While the molecular and neural mechanisms of circadian rhythms are well characterized, how cellular organization of clock components controls circadian clock regulation remains poorly understood. Here, we elucidate how clock proteins regulate circadian rhythms by controlling the spatiotemporal organization of clock genes. Using high-resolution live imaging techniques we demonstrate that *Drosophila* clock proteins are concentrated in a few discrete foci and are organized at the nuclear envelope; these results are in contrast to longstanding expectations that clock proteins are diffusely distributed in the nucleus. We also show that clock protein foci are highly dynamic and change in number, size, and localization over the circadian cycle. Further, we demonstrate that clock genes are positioned at the nuclear periphery by the clock proteins precisely during the circadian repression phase, suggesting that subnuclear localization of clock genes plays an important role in the control of rhythmic gene expression. Finally, we show that Lamin B receptor, a nuclear envelope protein, is required for peripheral localization of clock protein foci and clock genes and for normal circadian rhythms. These results reveal that clock proteins form dynamic nuclear foci and play a hitherto unexpected role in the subnuclear reorganization of clock genes to control circadian rhythms, identifying a novel mechanism of circadian regulation. Our results further suggest a new role for clock protein foci in the clustering of clock-regulated genes during the repression phase to control gene co-regulation and circadian rhythms.

**SIGNIFICANCE:** Almost all living organisms have evolved circadian clocks to tell time. Circadian clocks regulate ∼24-hour oscillations in gene expression, behavior and physiology. Here, we reveal the surprisingly sophisticated spatiotemporal organization of clock proteins and clock genes and its critical role in circadian clock function. We show, in contrast to current expectations, that clock proteins are concentrated in a few discrete, dynamic nuclear foci at the nuclear envelope during the repression phase. Further, we uncovered several unexpected features of clock protein foci, including their role in positioning the clock genes at the nuclear envelope precisely during the repression phase to enable circadian rhythms. These studies provide fundamental new insights into the cellular mechanisms of circadian rhythms and establish direct links between nuclear organization and circadian clocks.

## INTRODUCTION

Circadian clocks have a period of ∼24 hours and orchestrate daily rhythms in gene expression and affect many physiological processes ranging from sleep to metabolism to immunity (1). Circadian rhythms are self-sustained and can be entrained to environmental cues such as light(2) and temperature (3, 4). While past work has identified the molecular and neural basis of circadian clocks (5-9), the cellular mechanisms underlying circadian rhythms and how spatial organization of clock components control clock function remain largely unknown. Specifically, surprisingly little is known about clock protein subcellular localization and dynamics in live clock cells in vivo. Further, virtually nothing is known about how clock-regulated genes are spatially organized in individual clock cell nuclei in native tissues under physiological conditions, as past biochemical studies were performed on in vitro cell culture models (10, 11) or whole mammalian tissues (12, 13).

We address these critical gaps using *Drosophila melanogaster* which has a highly conserved yet relatively simple clock system consisting of 150 neurons (14) (Fig. 1a). Briefly, the *Drosophila* circadian clock consists of four key proteins—CLOCK (CLK), CYCLE (CYC), PERIOD (PER) and TIMELESS (TIM)—that form the core of the feedback loop and enable rhythmic activation and repression of the clock-regulated genes over the circadian cycle (15). Previous biochemical studies have shown that CLK and CYC proteins bind to the promoters of clock-regulated genes, including core clock genes-*per* and *tim*, and drive transcription during the activation phase (16) (Fig. 1b). Repression phase is initiated when PER and TIM proteins enter the nucleus, after a time delay, and inhibit CLK activity thus silencing their own expression as well as the expression of all other clock-regulated genes (17) (Fig. 1b). PER and TIM proteins are then degraded leading to the end of the repression phase and the start of a new cycle. A similar negative feedback loop functions in mammals with many of the clock proteins conserved among the two model systems (18). Remarkably, this molecular feedback loop operates in virtually every cell in the human body (19), from liver cells to neurons to skin cells, in order to ensure that cellular processes occur at the right time of the day and to generate circadian rhythms in metabolism, physiology, and behavior.

**Fig. 1:**
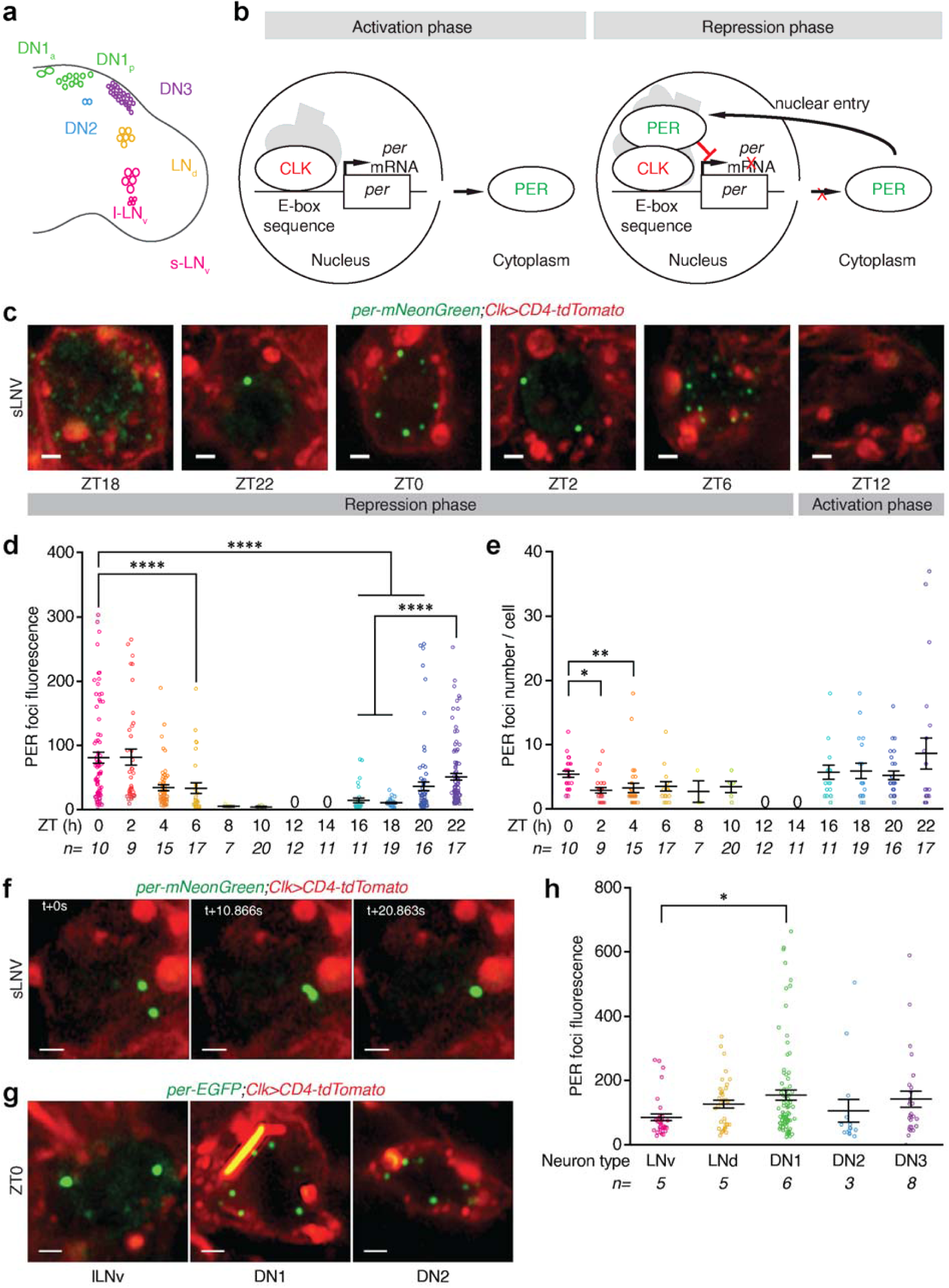
PER protein is concentrated in a few discrete nuclear foci during the circadian repression phase. (**a**) Schematic of the *Drosophila* circadian clock network, with the major classes of clock neurons labeled. (**b**) Schematic of the core molecular clock in the *Drosophila* clock neurons. During the activation phase, CLK protein complex binds to canonical E-box sequences and drive transcription of clock-controlled genes, including *period*. During the repression phase, PER protein enters the nucleus and inhibits CLK transcription factor activity silencing its own expression and all other clock-regulated genes. (**c**) Data from *per- mNeonGreen;Clk-Gal4>UAS-CD4-tdTomato* flies entrained to Light-Dark cycles (ZT0-lights on, ZT12-lights off). Representative images of PER foci (green) in a group of clock neurons, sLNVs (cell membrane labeled with tdTomato and shown in red), over the circadian cycle. (**d** and **e**) Quantitation of PER foci fluorescence (d) and foci number per sLNV (e) at specific ZTs over the 24-hour LD cycle. (**f**) Representative time-lapse images of sLNvs showing PER foci undergoing fusion. (**g**) Representative images of PER foci at ZT0 in other clock neurons (lLNvs, DN1s, and DN2s). (**h**) Quantitation of PER foci fluorescence in all classes of clock neurons at ZT0. Scale bars, 1 μm. Statistical tests used were a Kruskal-Wallis test (d, e, and h). \**P* < 0.05, \*\**P* < 0.005, \*\*\**P* < 0.0005, \*\*\*\**P* < 0.0001. Individual data points, mean, and s.e.m. (standard error of mean) are shown. ‘n’ refers to number of hemi-brains. See Supplementary Table 2 for detailed statistical analysis.

Clock-regulated genes, which number in hundreds and are spread throughout the genome (20), are co-regulated and rhythmically turned on and off once each day. It has also been shown that a large fraction of the mammalian genome is under circadian control (21). Our current understanding of circadian gene expression is based on the following model. All clock-regulated genes possess a canonical E-box sequence in their promoter region (16). These genes are activated by the binding of CLK protein and other transcriptional activators to their E-box sequences (16) (positive limb of the circadian feedback loop), and silenced when PER binds to CLK protein and recruits other transcriptional repressors to the chromatin (17, 22) (negative limb of the circadian feedback loop). However, we know virtually nothing about how all the clock-regulated genes are synchronously turned on and off once each day by the clock machinery. Further, while many of the advances in understanding the genetic and molecular mechanisms of circadian rhythms have come from either genetic screens or biochemical studies, very little is known about how spatial organization of clock components affect circadian clock function as it has not been possible so far to study the cellular organization and dynamics of clock proteins and clock genes in their native milieu.

Here, we investigate how cellular organization of clock proteins and clock genes in time and space control circadian clock regulation using *Drosophila* clock neurons as our model system. Towards this goal, we developed CRISPR-based single cell live imaging techniques and immuno-DNA fluorescent in-situ hybridization (FISH) approaches. Using these techniques, strikingly, we found that, in contrast to previous expectations (23-25), clock proteins are concentrated in a few discrete nuclear foci at the nuclear envelope. We also found that clock proteins position the clock genes at the nuclear periphery specifically during the repression phase when the genes are not transcriptionally active, suggesting that subnuclear reorganization of clock genes might play a key role in their rhythmic activation and repression. To elucidate how the nuclear envelope affects circadian rhythms, we conducted a genetic screen and found that loss of Lamin B receptor, a protein in the inner nuclear membrane, leads to disruption of clock protein foci and clock gene peripheral localization and results in circadian rhythm defects. These studies reveal a key mechanism of circadian regulation, which is likely broadly conserved, and suggest that clock protein foci regulate dynamic clustering and spatial reorganization of clock-regulated chromatin to control circadian rhythms in behavior and physiology.

## RESULTS

In order to elucidate the spatial distribution and dynamics of clock proteins in vivo at single-cell resolution, we generated fluorescent protein tagged flies in which endogenous PER was labeled with mNeonGreen (26). mNeongreen is a monomeric green/yellow fluorescent protein that is ∼3- to 5-fold brighter than EGFP and its maturation time is ∼3-fold shorter than that of EGFP (26, 27). Specifically, we applied CRISPR/Cas9 genome editing to fuse mNeongreen tag to the terminal exon of the endogenous *per* locus (Extended Data Fig. 1a, b), as previous mammalian studies have shown that PERIOD2-LUCIFERASE fusion protein is functional (19). To determine whether the PER-mNeonGreen fusion protein is functional in vivo, we analyzed whether *per* mRNA levels oscillate with a period of ∼24 hours. qPCR analysis revealed that *per* mRNA in these flies undergoes circadian oscillations and no significant differences were detected between wild-type (WT) and *per-mNeonGreen* flies (Extended Data Fig. 1c). Next, we analyzed the circadian behavior of *per-mNeonGreen* flies. These flies displayed WT-like behavior in both cycling (LD12:12) and constant conditions (DD), and they exhibited ∼24 hour free-running period rhythms, normal daily activity and sleep levels, and peaks of activity around dusk and dawn (Extended Data Fig. 2). Further, as shown in Extended Data Fig. 2, no significant circadian behavior differences were detected among homozygous *per-mNeonGreen* flies or heterozygous flies, demonstrating that PER-mNeongreen protein is functionally identical to the WT PER protein. Because a null mutation of the *per* locus causes loss of rhythmicity in constant conditions (28), we tested whether a single *per-*mNeonGreen allele can rescue the rhythms of *per^01^* null mutant flies. Female progeny from the cross (*per-mNeonGreen/per^01^*) have only one functional copy of *per-mNeonGreen* locus and can only produce PER-mNeonGreen fusion protein. We found that *per-mNeonGreen*/*per^01^* female flies displayed completely WT-like behavior in both cycling and constant conditions (Extended Data Fig. 2a, d), demonstrating that PER-mNeonGreen fusion protein is fully functional in vivo.

**Fig. 2:**
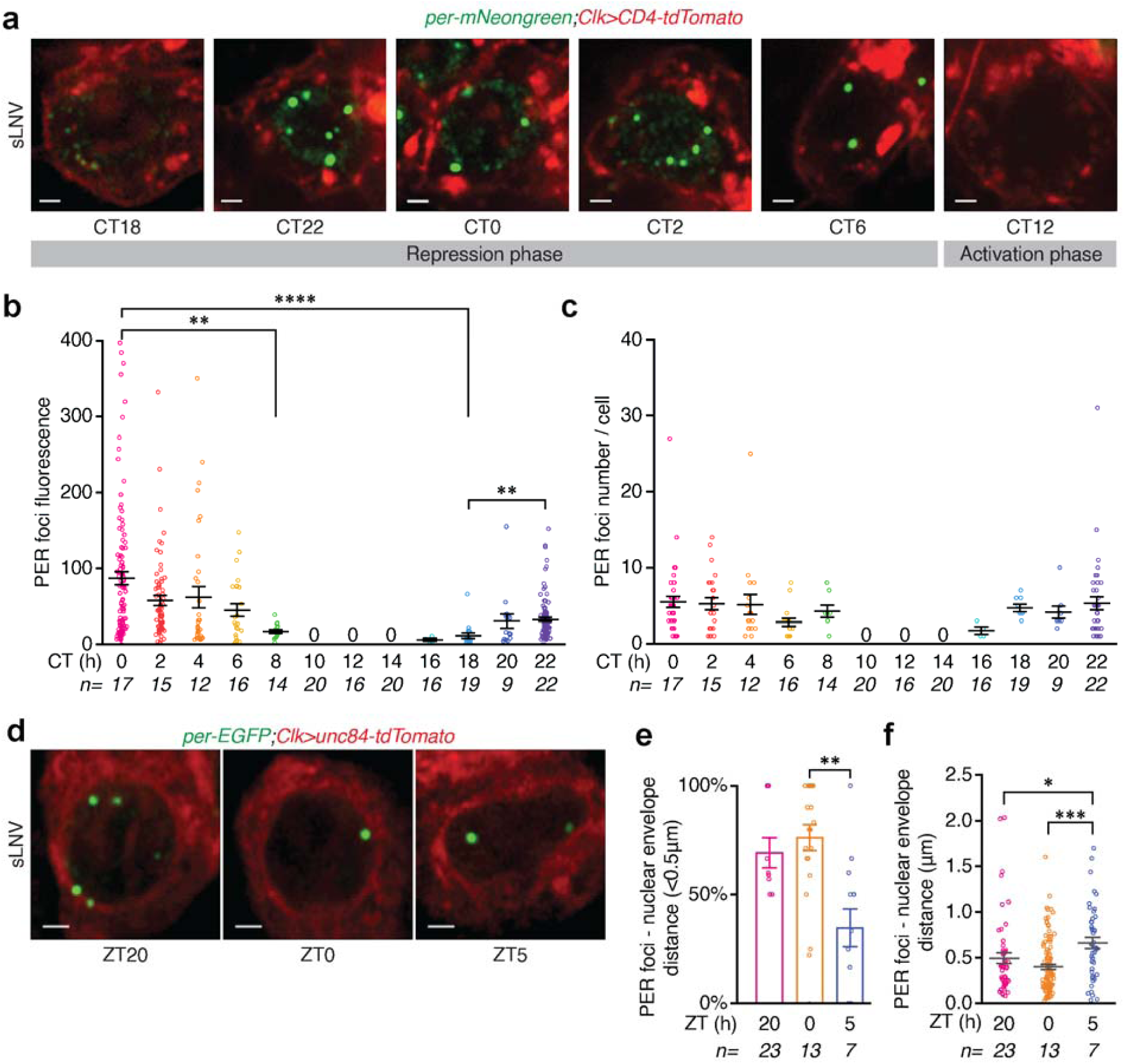
PER foci persist in constant conditions and are stereotypically positioned at the nuclear envelope. (**a**) *per-mNeonGreen;Clk-Gal4>UAS-CD4-tdTomato* flies are entrained to Light-Dark cycles (LD) for 5 days and released into complete darkness (DD). Representative images of PER foci in sLNVs at 2-hour intervals on day 1 of constant conditions in complete darkness. CT refers to ‘Circadian Time’. (**b** and **c**) Quantitation of PER fluorescence (b) and foci number per sLNV (c) at specific CTs during day 1 of constant conditions. (**d**) Representative images of PER foci in sLNvs from *per-EGFP;Clk-Gal4>UAS-unc84-tdTomato* flies. PER foci are located close to the nuclear envelope (marked in red) during the peak repression phase. (**e**) Quantitation of percentage of PER foci which are less than 0.5 μm away from the nuclear envelope at different ZTs. (**f**) Quantitation of the distance between PER foci and the nuclear envelope at different ZTs. Scale bars, 1 μm. Statistical tests used were a Kruskal-Wallis test (b, c, e, and f). \**P* < 0.05, \*\**P* < 0.005, \*\*\**P* < 0.0005, \*\*\*\**P* < 0.0001. Individual data points, mean, and s.e.m. (standard error of mean) are shown. ‘n’ refers to number of hemi-brains. See Supplementary Table 2 for detailed statistical analysis.

To monitor PER protein localization and dynamics in individual clock neurons in the brain, we entrained *per-mNeonGreen;Clk-Gal4>UAS-CD4-tdTomato* flies (*Clk-Gal4* is a pan clock neuron driver (29) and CD4 is a transmembrane protein that labels cell membranes (30)) to Light-Dark cycles (LD12:12, ZT0 - lights on, ZT12 - lights off) and imaged PER protein in individual clock neurons every 2 hours over the 24-hour cycle. We chose to first focus on small ventro-lateral neurons (sLNVs) which express PIGMENT DISPERSING FACTOR (PDF) neuropeptide, as they are considered to be the master pacemaker neurons (9, 31, 32). Our live imaging experiments performed using an Airyscan high-resolution confocal microscope revealed that PER protein is localized and is concentrated in discrete, dynamic foci in the nucleus of sLNVs during the repression phase of the circadian cycle (ZT18-ZT8) (Fig. 1c-e, Extended Data Fig. 3a, Supplementary mov. 1-3). These results are surprising and are in strong contrast to long-standing expectations that PER protein is diffusely distributed in the nucleoplasm (23, 24), which are based on past studies in clock neurons using static methods such as fixation and immunofluorescence. In fact, we observed that a majority of the PER foci disassemble upon formaldehyde fixation (Extended Data Fig. 3b), explaining why these foci escaped detection in previous studies. Another previous in vitro live imaging study (25) has shown that PER protein, when overexpressed in *Drosophila* S2 cells, forms cytoplasmic foci but shows diffuse localization in the nucleus. However, *Drosophila* S2 cells are regarded as “non-rhythmic” cells as they do not express several clock genes including *clk*, *per*, and *tim*, and it is unclear whether ectopic expression of clock genes can induce circadian rhythms. Moreover, overexpression of proteins can lead to aggregation artefacts. Our live imaging experiments using tagged endogenous proteins overcome all the limitations of past approaches and enables visualization of clock protein localization and dynamics at high spatio-temporal resolution in individual clock neurons within intact *Drosophila* brains.

**Fig. 3:**
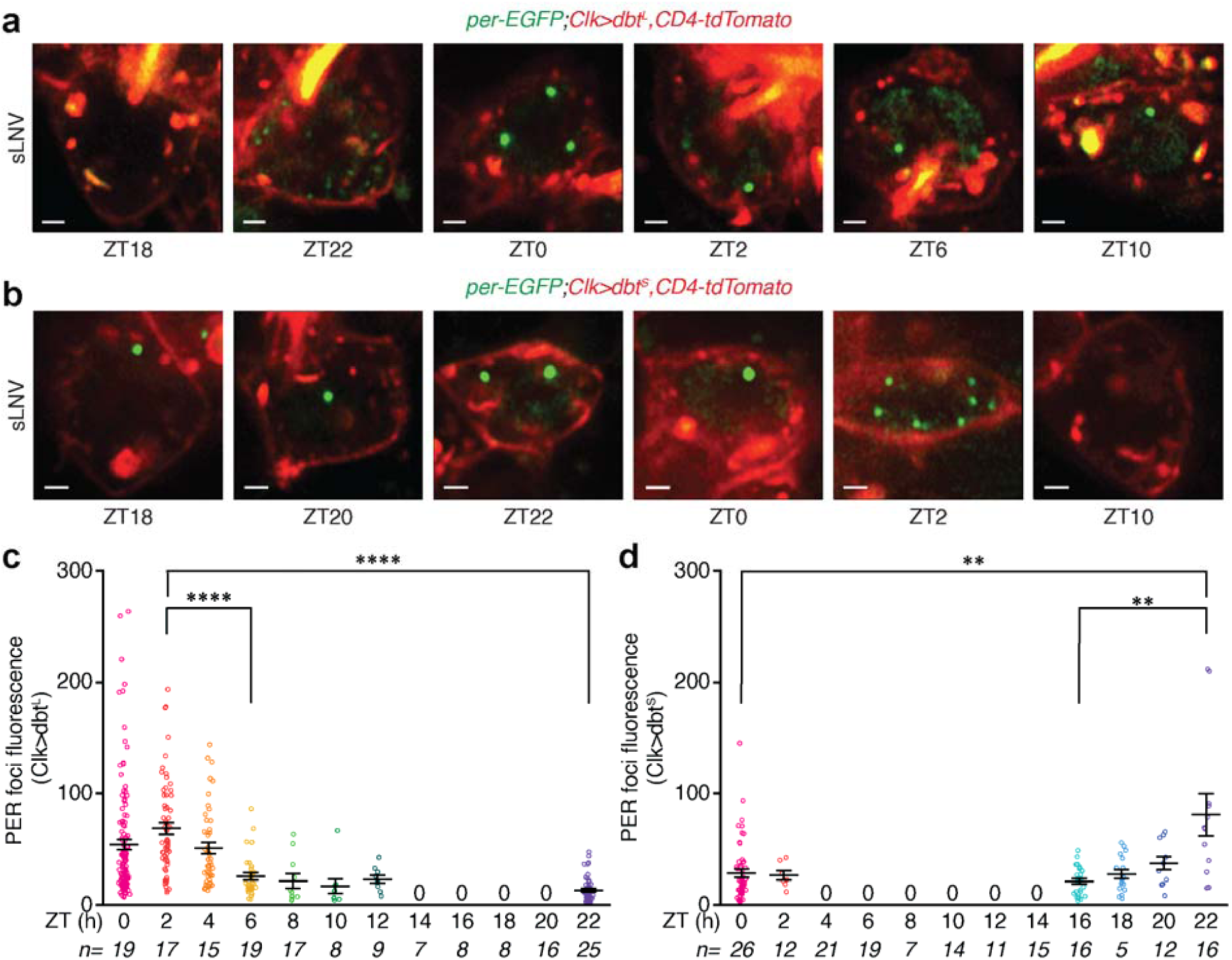
PER foci dynamics and spatial organization are regulated by Doubletime (DBT) kinase. (**a** and **b**) Representative images of sLNVs from *per-EGFP;Clk-Gal4>UAS-dbt^L^,UAS- CD4-tdTomato*flies (a) and *per-EGFP;Clk-Gal4>UAS-dbt^S^,UAS-CD4-tdTomato* flies (b) over the circadian cycle. (**c** and **d**) Quantitation of PER foci fluorescence in sLNvs from *dbt^L^* (c) and *dbt^S^* (d) mutants. Scale bars, 1 μm. Statistical tests used were a Kruskal-Wallis test (c, and d). \**P* < 0.05, \*\**P* < 0.005, \*\*\**P* < 0.0005, \*\*\*\**P* < 0.0001. Individual data points, mean, and s.e.m. are shown. ‘n’ refers to number of hemi-brains. See Supplementary Table 2 for detailed statistical analysis.

Specifically, our in vivo live imaging studies show that PER foci first appear in the cytoplasm at ZT16 (Extended Data Fig. 4a), start to accumulate in the nucleus from ZT18 (Fig. 1c), and persist till the end of the repression phase (ZT8). Recent studies have shown that intracellular condensate/foci assembly is often driven by weak multivalent interactions between proteins with intrinsically disordered regions (IDRs) and/or low-complexity sequences and nucleic acids (33). Interestingly, we found that PER protein has large regions of disorder in its C-terminal (Extended Data Fig. 4b). Further, we found that these PER foci are highly dynamic and exhibit liquid-like properties: these foci are spherical, two or more foci could fuse with each other to make a bigger foci (Fig. 1f, Supplementary mov. 4), and treatment of brains with 1,6-hexanediol, an aliphatic alcohol that disrupts weak hydrophobic interactions(33), disassembled all the PER foci in the clock neurons (Extended Data Fig. 4c, d).

**Fig 4:**
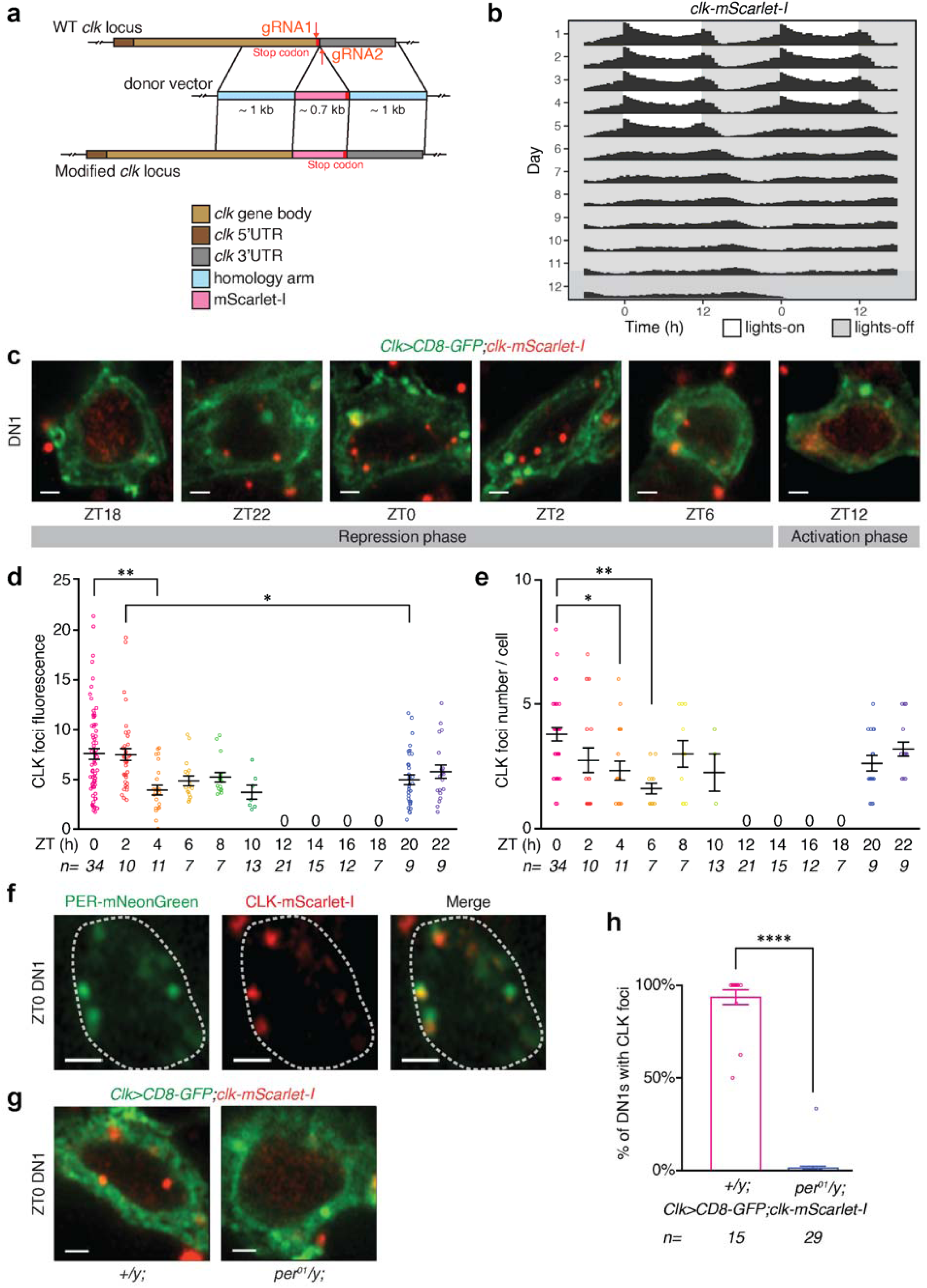
CLOCK forms nuclear foci in a PER-dependent fashion in the clock neurons during the repression phase. (**a**) Schematic of CRISPR/Cas9 genome editing to generate *clk-mScarlet-I* flies (See Methods for details). (**b**) *clk-mScarlet-I* flies were entrained to LD cycles (ZT0-lights on, ZT12-lights off) for 5 days and released into constant darkness for 7 days (DD). Averaged population locomotor activity profiles of *clk-mScarlet-I* flies (number of flies = 62) in LD and DD conditions with rest-activity shown for two consecutive days in the same line. These flies display rhythmic behaviors with a period of 24.39±0.10 hours with activity peaks around the time of lights-on and lights-off (See Extended Data Fig. 9b for details). (**c**) Representative images of CLK (red) in DN1s (cell membrane labeled with GFP and shown in green) from *Clk- Gal4>UAS-CD8-GFP;clk-mScarlet-I* flies over the circadian cycle. (**d** and **e**) Quantitation of CLK fluorescence (d) and foci number per DN1 (e) at specific ZTs over the 24-hour LD cycle. (**f**) Representative images of CLK and PER foci co-localization at ZT0 in DN1s from *per- mNeonGreen;clk-mScarlet-I* flies. 86% of CLK foci co-localize with PER foci (from 65 CLK foci from 10 hemi-brain images). (**g**) Representative images of CLK foci in DN1s at ZT0 from control (*+;Clk-Gal4>UAS-CD8-GFP;clk-mScarlet-I*) and *per^01^;Clk-Gal4>UAS-CD8-GFP;clk- mScarlet-I* mutant flies. (**h**) Quantitation of percentage of DN1s with CLK foci at ZT0 in control and *per^01^* mutant flies. Scale bars, 1 μm. Statistical tests used were a Kruskal-Wallis test (d and e) and a Mann-Whitney test (h). \**P* < 0.05, \*\**P* < 0.005, \*\*\**P* < 0.0005, \*\*\*\**P* < 0.0001. Individual data points, mean, and s.e.m. are shown. ‘n’ refers to number of hemi-brains. See Supplementary Table 2 for detailed statistical analysis.

The timing of PER foci nuclear entry (ZT18) corresponds to the start of the repression phase, and is consistent with previous ChIP studies that have shown that PER protein binds to DNA, via another clock protein CLK, at ZT18 to initiate repression (17). We found that these PER foci continue to increase in size until ZT0 (peak repression phase) and then decrease in size till PER protein is completely degraded around ZT8 (Fig. 1d). It is notable that PER protein foci are present long after (∼8 hours) lights are turned on at ZT0 (Fig. 1d, e), which has previously been shown to lead to rapid degradation of TIM protein (34), suggesting that PER protein persist as foci even after TIM is eliminated by light. These foci are present in almost all the sLNv clock neurons especially during the peak repression phase (Extended Data Fig. 5a). We observed on average ∼5-10 PER foci per sLNv at ZT0 and a lesser number at later time points (Fig. 1e). We also observed PER foci in all other clock neuron groups in the *Drosophila* brains, suggesting that PER nuclear foci appear synchronously throughout the clock network (Fig. 1g, h, Extended Data Fig. 5c). To further confirm our findings, we conducted live imaging studies using a different fly line where endogenous PER was labeled with an EGFP tag (35), which has been shown to display normal circadian rhythms. We found that PER-EGFP protein also forms discrete nuclear foci in clock neurons during the repression phase (Extended Data Fig. 6).

**Fig. 5:**
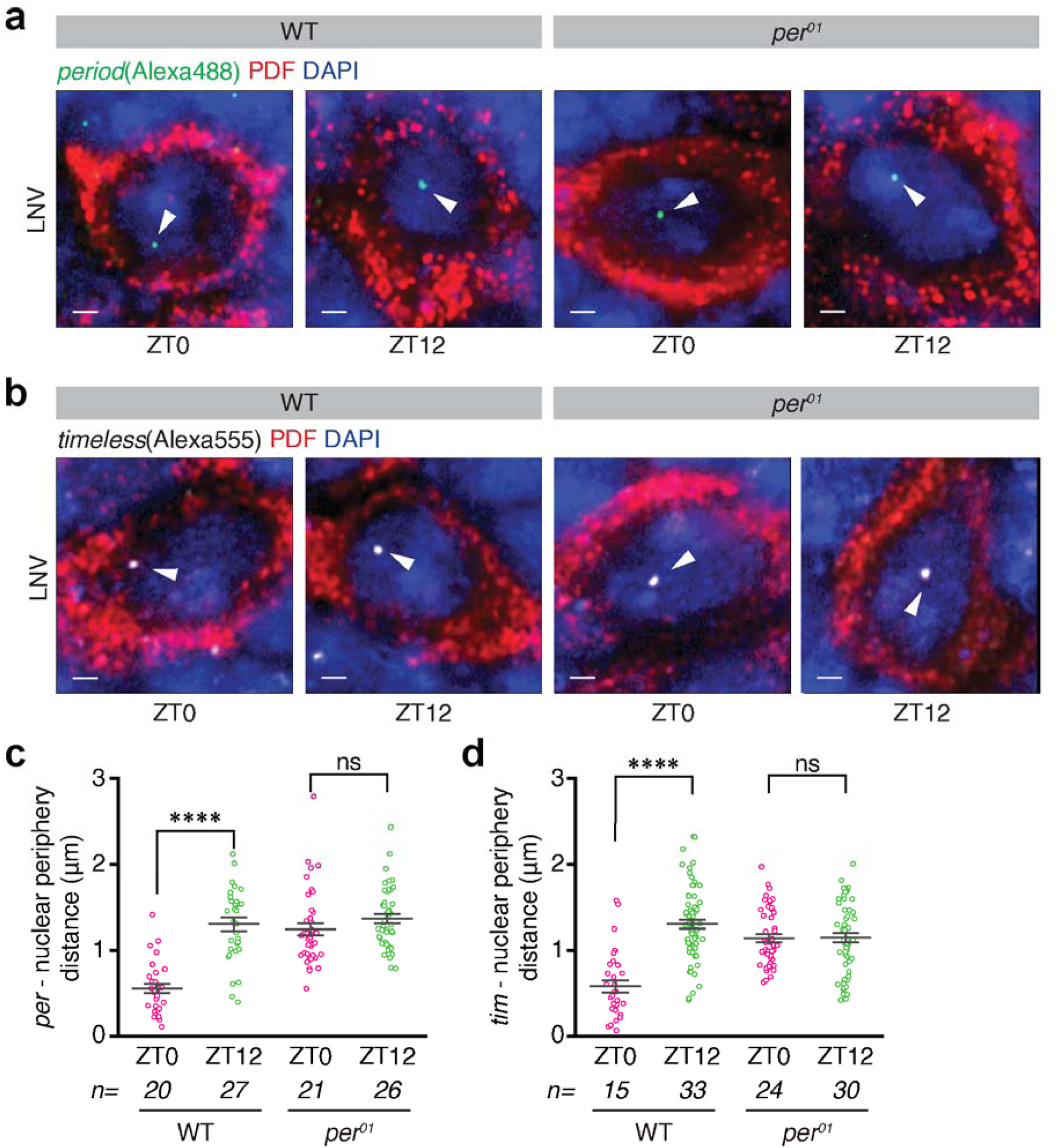
Core clock genes are positioned at the nuclear periphery by the clock proteins during the repression phase. (**a**) Representative images of DNA-FISH results using *period*- gene probes (green dots) in LNvs during repression (ZT0) and activation (ZT12) phases from wild-type (WT) and *per^01^* null mutant flies. Anti-PDF antibody (red) was used to identify LNv clock neurons and DAPI (blue) was used to mark the nucleus boundary. White arrowheads denote the location of *period* gene in the nucleus. (**b**) Representative images of DNA-FISH results using *timeless*-gene probes (white dots) in LNvs at ZT0 and ZT12 from wild-type and *per^01^* null mutant flies. White arrowheads denote the location of *timeless* gene in the nucleus. (**c** and **d**) Quantitation of distance between *period* gene and nucleus boundary (c) and *timeless* gene and nucleus boundary (d) in LNvs at ZT0 and ZT12 from wild-type and *per^01^* mutant flies. Scale bars, 1 μm. Statistical tests used were a Kruskal-Wallis test (c and d). \**P* < 0.05, \*\**P* < 0.005,\*\*\**P* < 0.0005, \*\*\*\**P* < 0.0001. Individual data points, mean, and s.e.m. are shown. ‘n’ refers to number of hemi-brains. See Supplementary Table 2 for detailed statistical analysis.

**Fig. 6:**
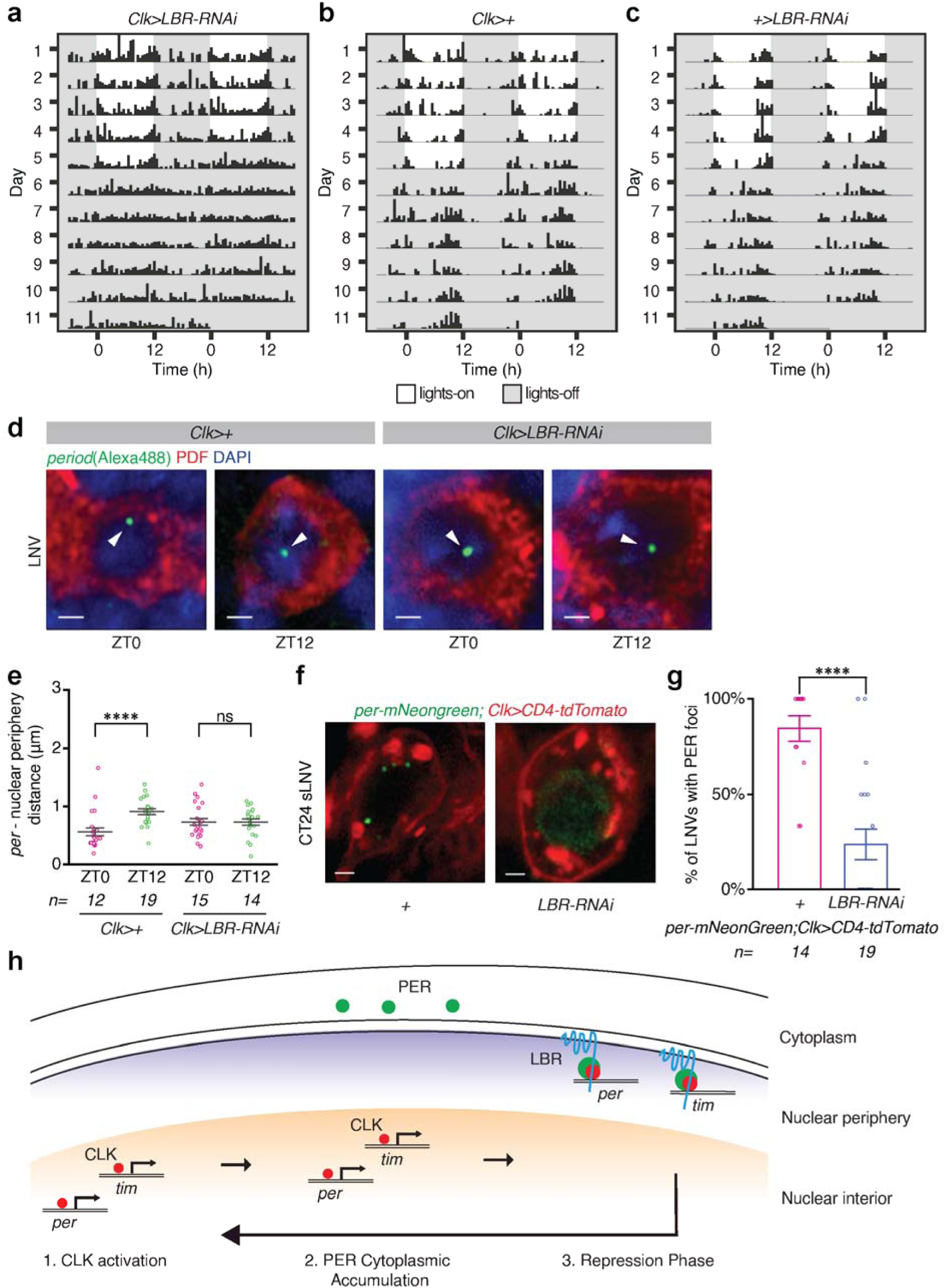
Lamin B Receptor (LBR) tethers clock genes to the nuclear envelope and regulates circadian rhythms. (**a**, **b** and **c**) Representative activity profile of individual *Clk-Gal4>UAS- LBR-RNAi* (a), *Clk-Gal4>+*(b), +*>UAS-LBR-RNAi* (c) flies with rest-activity shown for two consecutive days in the same line. (See Extended Data Fig. 10 for details). (**d**) Representative images of DNA-FISH results using *period*-gene probes (green dots) in LNvs at ZT0 and ZT12 from control (*Clk-Gal4>+*) and *Clk-Gal4>UAS-LBR-RNAi* flies. (**e**) Quantitation of distance between *period* gene and nucleus boundary in LNvs at ZT0 and ZT12 from control (*Clk- Gal4>+*) and *Clk-Gal4>UAS-LBR-RNAi* flies. **(f)** Representati ve images of PER foci in sLNVs in control (*per-mNeonGreen;Clk-Gal4>UAS-CD4-tdTomato,+*) and experimental (*per- mNeonGreen;Clk-Gal4>UAS-CD4-tdTomato,LBR-RNAi*) flies at CT24 on the second day of constant conditions. **(g)** Quantitation of percentage of sLNvs with PER foci in control and experimental flies at CT24. (**h**) A model for the spatiotemporal localization of clock protein- chromatin complexes in clock neurons over the circadian cycle. Scale bars, 1 μm. Statistical tests used were a Kruskal-Wallis test (e and g). \**P* < 0.05, \*\**P* < 0.005, \*\*\**P* < 0.0005, \*\*\*\**P* < 0.0001. Individual data points, mean, and s.e.m. are shown. ‘n’ refers to number of hemi-brains. See Supplementary Table 2 for detailed statistical analysis.

To test whether PER-mNeonGreen foci persist in constant darkness, we entrained the flies to LD12:12 cycles and released them into constant darkness and imaged PER protein in the first day of constant conditions (DD). We found that PER foci persist in clock neuron nuclei in constant darkness after entrainment and the number and size of PER foci is not significantly different in LD and DD conditions (Fig. 2a-c, Extended Data Fig. 5b), suggesting that PER protein oscillations do not dampen immediately upon transfer to constant conditions. Notably, we observed PER foci in clock neurons even after 8 days in complete darkness (Extended Data Fig. 5d, e). In addition to oscillations in PER foci size and number, we found that PER foci also show a stereotypical subnuclear localization at the nuclear envelope during the repression phase of the circadian cycle. To visualize the nuclear envelope clearly and quantify the distance between PER foci and the nuclear envelop precisely, we crossed *per-mNeonGreen* flies to *Clk- Gal4;UAS-unc84-tdTomato* flies (UNC84 is a SUN domain protein that localizes to the inner nuclear membrane (36)) and imaged PER foci during the repression phase. We observed that a majority of the PER foci are located close to the nuclear envelope during the repression phase (Fig. 2d-f, Supplementary mov. 5-7). Furthermore, PER foci at ZT0 (peak repression phase) exhibited significantly less mean squared displacement compared to the foci at ZT5 (Extended Data Fig. 7), suggesting that PER foci at ZT0 might be spatially constrained at the nuclear envelope. Together, these studies establish that PER protein forms dynamic nuclear foci in clock neurons that oscillate with a 24-hour rhythm in their size, number, and subcellular localization.

Next, we examined PER foci dynamics in clock mutants in which the 24-hour period of circadian rhythms is either shortened or lengthened. Specifically, we examined mutants of a key clock kinase, Casein Kinase I/DOUBLETIME (DBT), which is known to phosphorylate PER protein and regulate its stability and activity (37). Flies that ectopically express *dbt^L^* mutation (*UAS-dbt^L^*) specifically in clock neurons display circadian rhythms with an average free-running period of ∼27 hours (38) (Extended Data Fig. 8a, b), while *dbt^S^* mutation in clock neurons produces rhythms with an average free-running period of 18 hours (38) (Extended Data Fig. 8a, c). In our experiments using *Clk-Gal4>UAS-dbt^L^* flies, we found that PER foci start to appear in the nucleus at ZT22 and persist until ZT12 (Fig. 3a, c, Extended Data Fig. 8d), which is later compared to WT conditions, suggesting that *dbt^L^* mutation causes delayed nuclear entry. In contrast, in *Clk-Gal4>UAS-dbt^S^* flies PER foci appear in the nucleus at ZT16, which is slightly earlier compared to WT, and these foci disappear by ZT4 (Fig. 3b, d, Extended Data Fig. 8e), suggesting that PER protein is prematurely degraded in the *dbt^S^* mutant flies. These results show that PER foci dynamics and spatial distribution are regulated by DBT kinase, indicating that the timing of PER foci nuclear entry and their degradation affect circadian period length.

Previous biochemical studies have shown that PER initiates transcriptional repression by binding to CLK protein, a positive regulator of the circadian feedback loop, which is bound to E-box elements of the clock-regulated genes (17, 22). To visualize CLK protein, we generated fluorescent protein tagged flies in which endogenous CLK was labeled with mScarlet-I(39), a bright monomeric red fluorescent protein (Fig. 4a, Extended Data Fig. 9a). We established that the tagged protein (CLK-mScarlet-I) is functional by showing that the behavior of the homozygous *clk-mScarlet-I* flies and heterozygous flies is rhythmic (Fig. 4b, Extended Data Fig. 9b-d), and *per* mRNA in these flies undergoes circadian oscillations similar to wild-type levels (Extended Data Fig. 1c). To perform live imaging experiments, we entrained *Clk-Gal4>UAS- CD8-GFP;clk-mscarlet-I* flies (CD8 is a transmembrane protein that labels cell membranes(30)) to LD cycles (LD12:12, ZT0 - lights on, ZT12 - lights off) and imaged CLK protein every 2 hours.

We found that CLK protein is also concentrated in discrete nuclear foci in the clock neurons specifically during the repression phase of the circadian cycle (Fig. 4c-e). These results are in contrast to previous observations that CLK protein is diffusely located in the nucleus based on immunostaining studies on fixed brains (40). Importantly, we did not detect any distinct nuclear CLK foci in the clock neurons during the activation phase (Fig. 4c-e), strongly suggesting the CLK foci are formed specifically during the repression phase. CLK nuclear foci start to appear at ZT20, ∼2 hours after PER protein enters the nucleus, increase in size until peak repression phase (ZT0) and persist till the end of the repression phase (∼ZT8) (Fig. 4d, e). We also found that CLK foci co-localize with the PER foci during the circadian repression phase (Fig. 4f). We note that we also found a few red foci (CLK protein) outside clock neurons; these observations are consistent with previous studies which have shown that CLK protein is found in many glial cells as well as other non-clock neurons in the brain (40, 41). Next, we tested whether PER is necessary for CLK foci formation in the clock neurons by examining *per^01^* null mutant flies. Notably, we did not find any CLK foci in the clock neuron nuclei of *per^01^* mutant flies; instead CLK is diffusely distributed in the nucleus in the absence of PER (Fig. 4g, h), demonstrating that PER is required for CLK protein to be aggregated into discrete foci. From the above studies, we conclude that CLK protein is diffusely located in the clock neuron nuclei during the activation phase when it drives transcription of circadian genes, and CLK protein is organized into discrete nuclear foci by the PER protein during the repression phase of the circadian cycle.

Next, we examined how PER-CLK foci may inhibit expression of circadian genes during the repression phase. Since past work has shown that PER-CLK complex binds to the promoters of clock-regulated genes during the repression phase (17, 22), and that repressed chromatin is generally located at the nuclear periphery (42, 43), we hypothesized that circadian genes might also be rhythmically positioned at the nuclear envelope by the clock proteins during the repression phase. To test our hypothesis, we adapted an in situ immuno-DNA FISH technique for use in adult *Drosophila* brains (see Methods). Our DNA-FISH experiments show that core clock genes, *per* and *tim*, are positioned at the nuclear periphery in clock neurons specifically during the repression phase (ZT0) and are located in the nuclear interior during the activation phase (ZT12) (Fig. 5a-d). Importantly, nuclear peripheral positioning of *per* and *tim* genes is disrupted in *per^01^* mutants (Fig. 5a-d), suggesting that PER protein plays a critical role in the spatial organization of clock genes over the circadian cycle. We observed only one focus or two closely spaced foci per gene in all the clock neurons (Fig. 5a, b), consistent with the data that homologous chromosomes are paired in *Drosophila* somatic cells (44).

We note that we were not able to detect colocalization of PER-CLK protein foci and *per/tim* gene foci in clock neurons due to technical limitations, as the DNA-FISH procedure which includes the fixation process led to the disassembly of the clock protein foci (Extended Data Fig. 3b). However, this does not pose a challenge in interpreting our results as past studies using ChIP and proteomics techniques have unambiguously shown that PER protein binds to the promoters of the clock-regulated genes, via CLK, and recruits chromatin repressive complexes to enable gene silencing during the repression phase (17, 22). Based on the above results, we conclude that core clock genes in the clock neurons are positioned at the nuclear periphery by the clock proteins during the repression phase, and are repositioned to the nuclear interior during the activation phase. These results suggest that dynamic repositioning of clock genes to different subnuclear locations over the circadian cycle can control their rhythmic transcriptional activation and repression.

To determine the molecular mechanisms underlying the localization of clock protein foci and clock genes at the nuclear envelope, we conducted a behavioral screen of all known *Drosophila* lamin and nuclear envelope proteins (45, 46): *Lamin B*, *Lamin C*, *Lamin B Receptor* (*LBR*), *fs(1)Ya*, SUN-KASH domain proteins-*koi*, *klar*, *Msp300*, and LEM domain proteins- *Otefin*, *dMAN1*, *Bocksbeutel*. Specifically, we knocked down the expression of these genes in the clock neuron network using the *Clk-Gal4* driver. Through our screen, we identified that Lamin B receptor (LBR), an inner nuclear membrane protein that binds to both lamin and chromatin (47, 48), is required for locomotor activity rhythms (Extended Data Fig. 10a). LBR contains a nucleoplasmic N-terminal domain and a hydrophobic C-terminal consisting of eight transmembrane domains (47, 48). Previous studies have shown that LBR associates with chromatin via HP1 (49, 50) and it is required for peripheral heterochromatin organization in many cell types (51-53). In our behavior studies, we observed that knockdown of LBR expression in the clock neuron network (*Clk-Gal4>UAS-LBR-RNAi*) led to defects in rhythmic behavior compared to the parental controls (*Clk-Gal4>+*, *+>UAS-LBR-RNAi*) (Fig. 6a-c, Extended Data Fig. 10). Specifically, a majority of the experimental flies (63.3%) displayed arrhythmic behavior in constant conditions with no distinct activity peaks around subjective dawn and dusk (Extended Data Fig. 10a). Coincidentally, a past study (54) using a different *UAS-LBR-RNAi^KK110508^* line has also reported loss of behavioral rhythmicity, further strengthening our findings. We note that knockdown of LBR in the clock neurons did not affect the total number of clock neurons or the overall morphology of the clock neuron network.

To determine how LBR affects circadian rhythms, we tested whether disruption of LBR affects clock gene subnuclear localization. We performed DNA-FISH experiments and found that knockdown of LBR leads to defects in the localization of the *period* gene at the nuclear periphery during the circadian repression phase (Fig. 6d, e). Finally, we performed PER foci live imaging experiments on flies in which LBR expression is knocked down in all the clock neurons (*per-EGFP;Clk-Gal4>UAS-CD4-tdTomato,LBR-RNAi*). We observed that PER protein is diffusely located throughout the nucleus in a majority of clock neurons in the *clk>LBR-RNAi* flies compared to the control (*clk>+*) flies (Fig. 6f, g), suggesting that LBR might be required for PER protein to be concentrated in discrete foci during the repression phase. Taken together, our results suggest that LBR plays a key role in the regulation of circadian rhythms by controlling the peripheral positioning of clock genes and clock protein foci during the repression phase. These studies provide novel mechanistic insights into how the nuclear envelope regulates circadian gene expression at a global scale and controls circadian rhythms.

## DISCUSSION

Our findings reveal how the spatiotemporal organization of clock proteins and genes controls circadian clock regulation, providing critical new insights into how clocks function at the single-cell level. Our work demonstrates that clock proteins are concentrated in dynamic nuclear foci organized at the nuclear envelope, and play a crucial role in the repositioning of clock genes to the nuclear periphery during the circadian repression phase (Fig. 6h), suggesting a novel mechanism by which circadian clocks are regulated.

In our live imaging studies, surprisingly, we found that PER protein is concentrated in a few discrete foci located close to the nuclear envelope during the circadian repression phase (Fig. 1c), and is not diffusely located in the nucleus as expected from past studies (23-25). CLK protein, which is a positive transcription factor, is also found to be concentrated in nuclear foci that colocalize with PER foci during the repression phase, but is diffusely distributed in the nucleus during the activation phase (Fig. 4c). To test whether any of the *Drosophila* clock proteins possess intrinsically disordered regions, we analyzed the amino acid sequence of the proteins and found that PER protein has intrinsically disordered regions in its C-terminal (Extended Data Fig. 4b). Interestingly, recent studies have shown that clock proteins from many species contain intrinsically disordered regions (55-57), however, little is known about how disorder in clock proteins affects circadian clock function. As recent studies have shown that disordered regions promote phase separation and foci formation in other biological contexts(33), our results, which reveal that clock proteins spontaneously form foci at the nuclear envelope under physiological conditions, provide clues into the potential biological significance of the clock protein disordered regions in circadian rhythms.

Our work highlights multiple properties of the clock protein foci that enable circadian clock regulation, which we were able to uncover by studying the spatiotemporal organization and dynamics of clock proteins in their native milieu. First, we found that PER foci are highly dynamic and exhibit liquid-like properties (Fig. 1f). For example, we observed a number of fusion events especially at time points earlier than ZT0 (peak repression phase), in accordance with our observation that foci decrease in number and increase in size during the early repression phase (Fig. 1d, e). Second, we observed a gradual decrease in both the size and the number of PER foci at later times during the repression phase (Fig. 1d, e), which is consistent with the fact that PER protein is gradually degraded over the repression phase (34). Interestingly, these results point to a new hypothesis that PER foci might be heterogeneous and each individual PER focus might be regulated (*e.g.,* time of degradation) somewhat independently through yet unknown mechanisms. We favor the idea that PER foci might have different sets of DNA, RNA, and protein molecules that can be involved in differential regulatory processes, which can contribute to phase differences in the expression of clock-regulated genes within a single cell.

Third, we observed only a few PER foci (<10) per neuron during the circadian repression phase (Fig. 1d). Given that clock-regulated genes number in hundreds (21), these results raise the intriguing possibility that clock proteins might drive clustering of clock-regulated genes into a few nuclear foci during the repression phase to enable transcriptional co-regulation through common cis-acting elements (E-boxes). Past studies, which employed chromosome-conformation capture approaches (12, 13), have shown that enhancer-promoter interactions of core clock genes are under circadian control. However, how clock-regulated genes are spatially organized and clustered in the nuclear space and whether that leads to their co-regulation remains unknown. Based on our imaging and DNA-FISH results, we propose that clock-regulated genes might be clustered via inter-/intra- chromosomal interactions by the clock-protein complexes specifically during the repression phase, analogous to the compaction and clustering of repressed genes into discrete nuclear foci by Polycomb complexes (58, 59), referred to as ‘PcG bodies’ or ‘PcG clusters’.

Fourth, we show that clock proteins are required for positioning the clock genes to different subnuclear locations—to the nuclear periphery during repression phase and to the nuclear interior during the activation phase—over the circadian cycle (Fig. 5). To determine the underlying mechanisms linking the nuclear envelope to circadian rhythms, we leveraged the power of *Drosophila* genetics and conducted a genetic screen and identified that Lamin B receptor, located in the inner nuclear membrane, is required for peripheral localization of clock protein foci and clock genes during the repression phase and for circadian rhythms (Fig. 6). Interestingly, previous studies have also shown that LBR is required for peripheral heterochromatin organization for the silencing of one of the X-chromosomes in female mammalian cells (53).

Our work demonstrates that clock proteins form nuclear foci and have a novel function of positioning the clock genes to the nuclear envelope for regulating circadian rhythms. Past work on gene regulation has shown that spatial organization of the genome plays a crucial role in regulating gene expression, e.g., inactive X chromosome in female mammalian cells (60), immunoglobulin genes in hematopoietic stem cells (61), and stem cell differentiation genes (62). While these past studies point to a cell type-specific positioning of certain gene loci that is established at a specific stage during development, our studies indicate that clock-regulated genes are positioned to different subnuclear environments rhythmically every single day throughout the life of the organism. Therefore, we propose that circadian genome organization in post-mitotic clock neurons is highly dynamic in space and time, and chromatin movement and positioning to specific subnuclear locations might be critical for rhythmic circadian gene expression. How clock protein foci control positioning of clock genes to specific subnuclear locations at particular times over the circadian cycle and how chromosome dynamics affects circadian gene expression remains to be determined. Given the remarkable similarity between *Drosophila* and mammalian clock systems, we expect the cellular mechanisms uncovered here will be broadly applicable to circadian clocks in humans.

## ACKNOWLEDGEMENTS

The authors acknowledge fruitful discussions with Yukiko Yamashita, Orie Shafer, Josie Clowney, Edgar Meyhofer, and Pramod Reddy. The authors thank Yong Zhang, Orie Shafer, Paul Hardin, Henry Gilbert, Amita Sehgal, Justin Blau, Scott Pletcher, the Bloomington *Drosophila* Stock Center, the Developmental Studies Hybridoma Bank, and Addgene for fly stocks and other reagents. We also thank Pierre Coulombe and Scott Pletcher for sharing equipment in the initial stages of the project.

## Funding

The work was supported by a grant from the NIH (grant no. R35GM133737 to S.Y.) and the University of Michigan startup funds to S.Y..

## Author contributions

S.Y. conceived and designed the research; Y.X. designed the CRISPR experiments and constructed the transgenic flies under the supervision of S.Y.; Y.X. and Y.Y. designed the DNA FISH probes under the supervision of S.Y.; Y.X., Y.Y., and S.Y. conducted the live-imaging experiments, behavior experiments, qPCR experiments, and DNA FISH experiments; M.J. and S.Y. collected data for the initial per-EGFP imaging experiments; N.S helped in collecting data for the experiments presented in Fig. 2; Y.Y. developed data analysis scripts; Y.X., Y.Y., and S.Y. analyzed the data from all the experiments; S.Y. supervised the project; and Y.X., Y.Y., and S.Y. interpreted the results and wrote the manuscript.

## Competing interests

The authors declare no competing interests.

## Data availability

All of the data pertaining to the work are available in the manuscript or the supplementary materials. All the data analysis scripts from this study are available from the corresponding author upon reasonable request.

---

**Extended Data Fig. 1:**
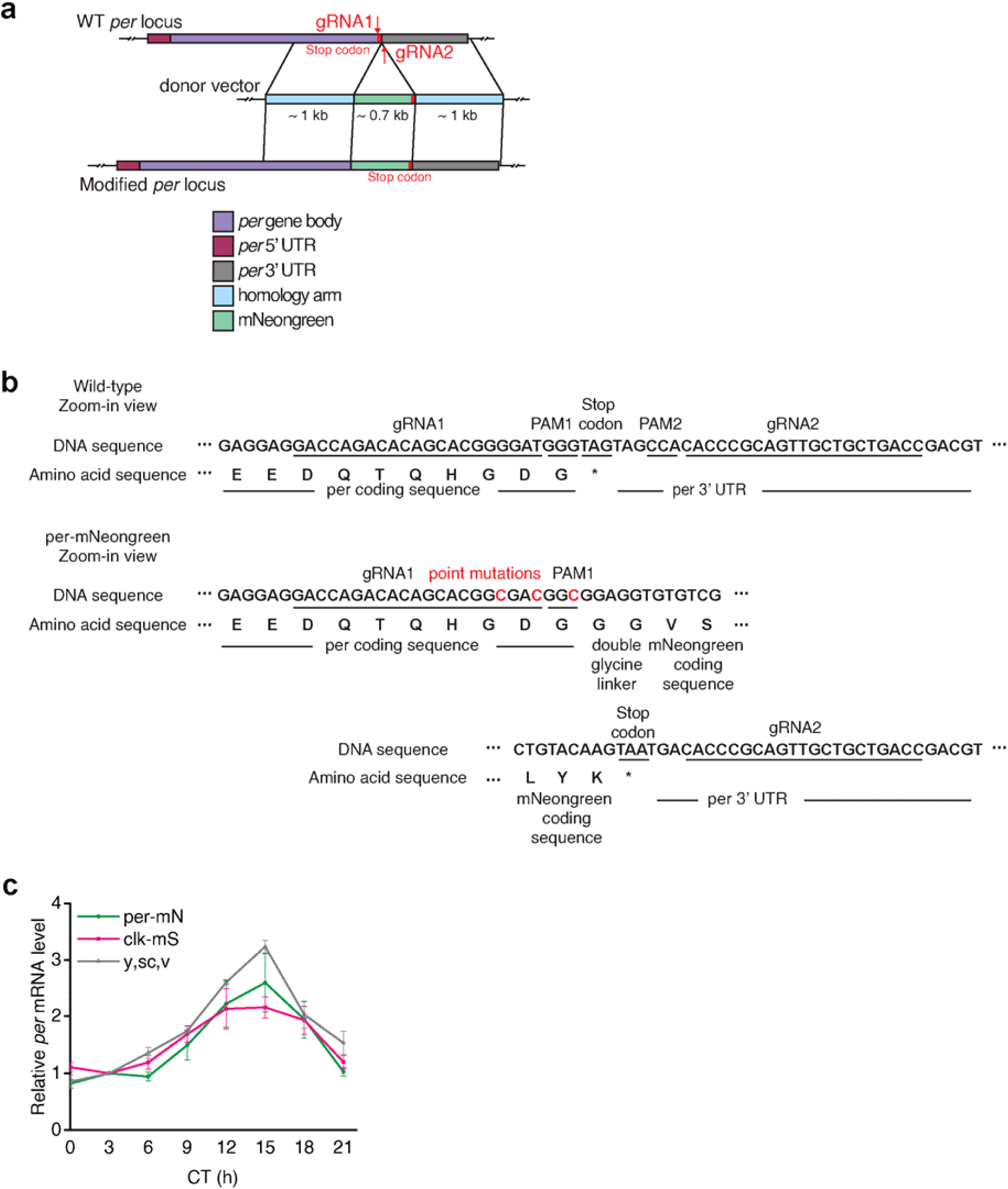
Generation and characterization of *per-mNeonGreen* fly. (**a**) Strategy for tagging the endogenous *period* gene with the mNeonGreen fluorescent tag. The two gRNAs are located on either side of the stop codon, one in the last exon and another in the 3’UTR sequence. The donor vector contains two ∼1 kb homology arms and mNeonGreen tag with stop codon attached to it. (**b**) Sequencing results of *period*-mNeonGreen allele. The protospacer adjacent motifs (PAM) of gRNA1 was mutated (amino acid sequence of the protein was not changed) to prevent cutting by Cas9 after homology directed repair events. Point mutations are marked in red. (**c**) Total mRNA extracted from ∼30 fly heads at different times of the first day in complete darkness (CT) was analyzed by qPCR. *period* mRNA levels were normalized to that of *rp49*. Data were averaged from three independent experiments and error bars represent s.e.m.. CT denotes ‘Circadian Time’ in complete darkness conditions after entrainment to LD cycles, and CT0 is the start of the subjective day and CT12 is the start of the subjective night. See Supplementary Table 2 for detailed statistical analysis.

**Extended Data Fig. 2:**
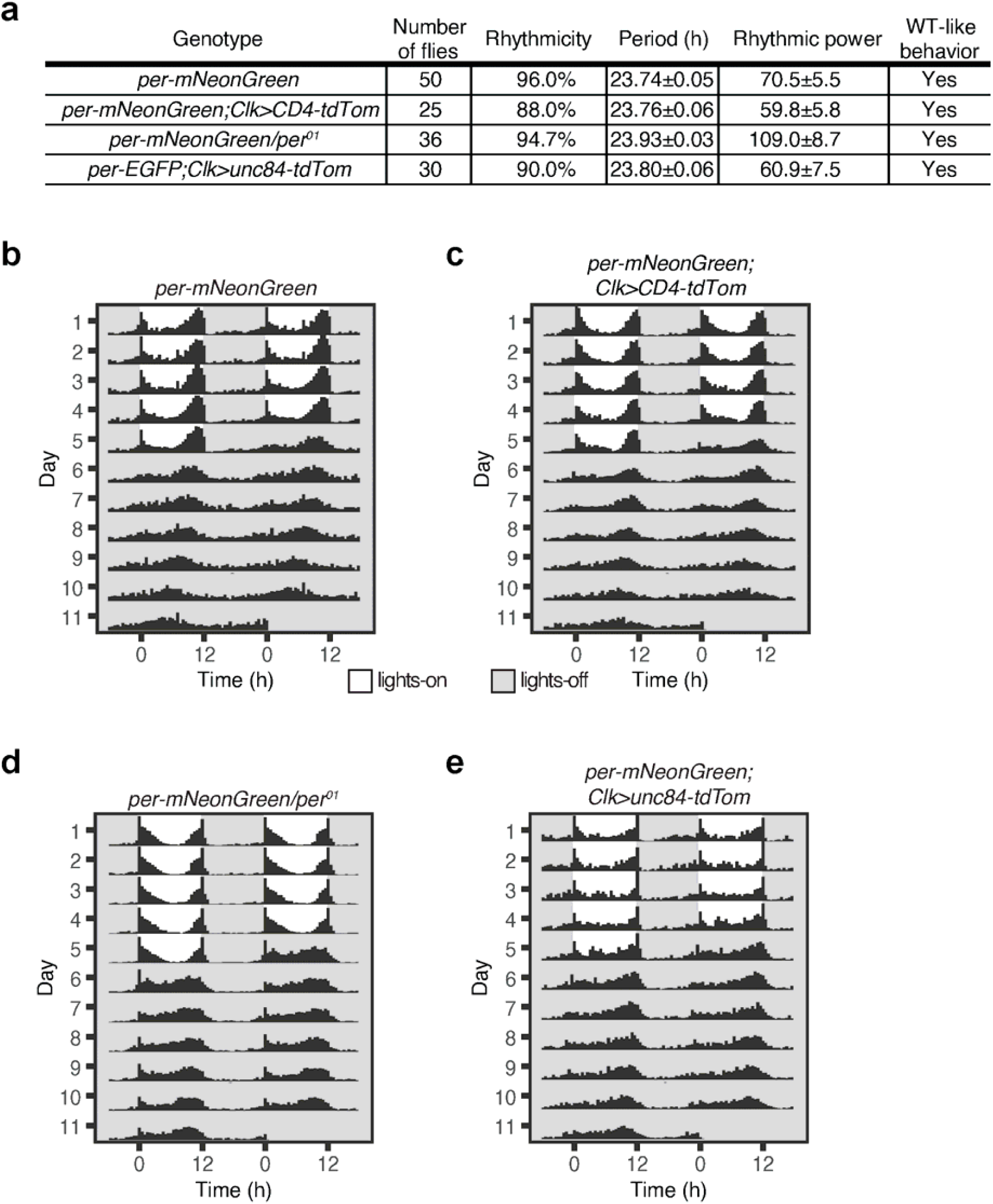
Behavioral rhythms of *per-mNeonGreen* flies. (**a**) Summary of free- running locomotor activity rhythms under constant conditions after entrainment to Light-Dark (LD) cycles. The genotypes of the flies tested are listed in the table. Period was calculated by chi-square periodogram analysis for all the flies and shown as average period ± s.e.m.. (**b**, **c**, **d, e**) Flies of indicated genotypes were entrained to LD cycles (ZT0-lights on, ZT12-lights off) for 5 days and released into constant darkness for 7 days. Averaged population locomotor activity profiles of flies with rest-activity shown for two consecutive days in the same line. These flies display rhythmic behaviors with activity peaks around the time of lights-on and lights-off

**Extended Data Fig. 3:**
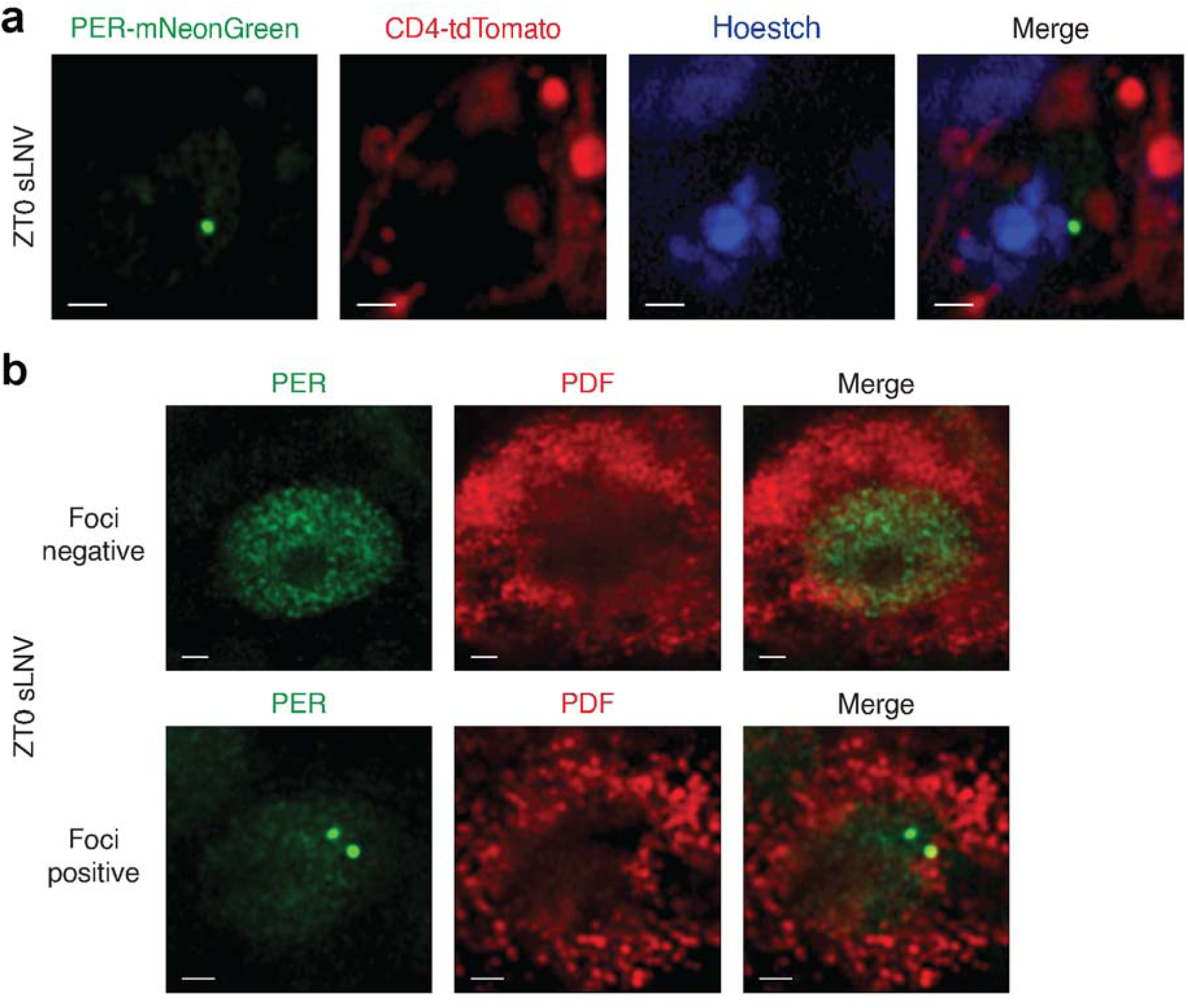
PER foci are located in the nucleus during the repression phase. (**a**) Representative images of PER foci (green) in sLNVs (cell membrane labeled with tdTomato and shown in red) at ZT0 from *per-mNeonGreen;clk>CD4-tdTomato* flies. Hoestch stain is used to stain DNA. PER foci are present in the nucleus of clock neurons during the repression phase. (**b**) Representative images of PER (green) in LNvs in brains, which were fixed with 4% paraformaldehyde and stained with Anti-PDF (red) antibodies. Majority of the clock neurons in fixed brains display diffuse PER staining, suggesting that PER foci are disassembled upon fixation. A very small fraction of cells after fixation still display PER foci (<5%).

**Extended Data Fig. 4:**
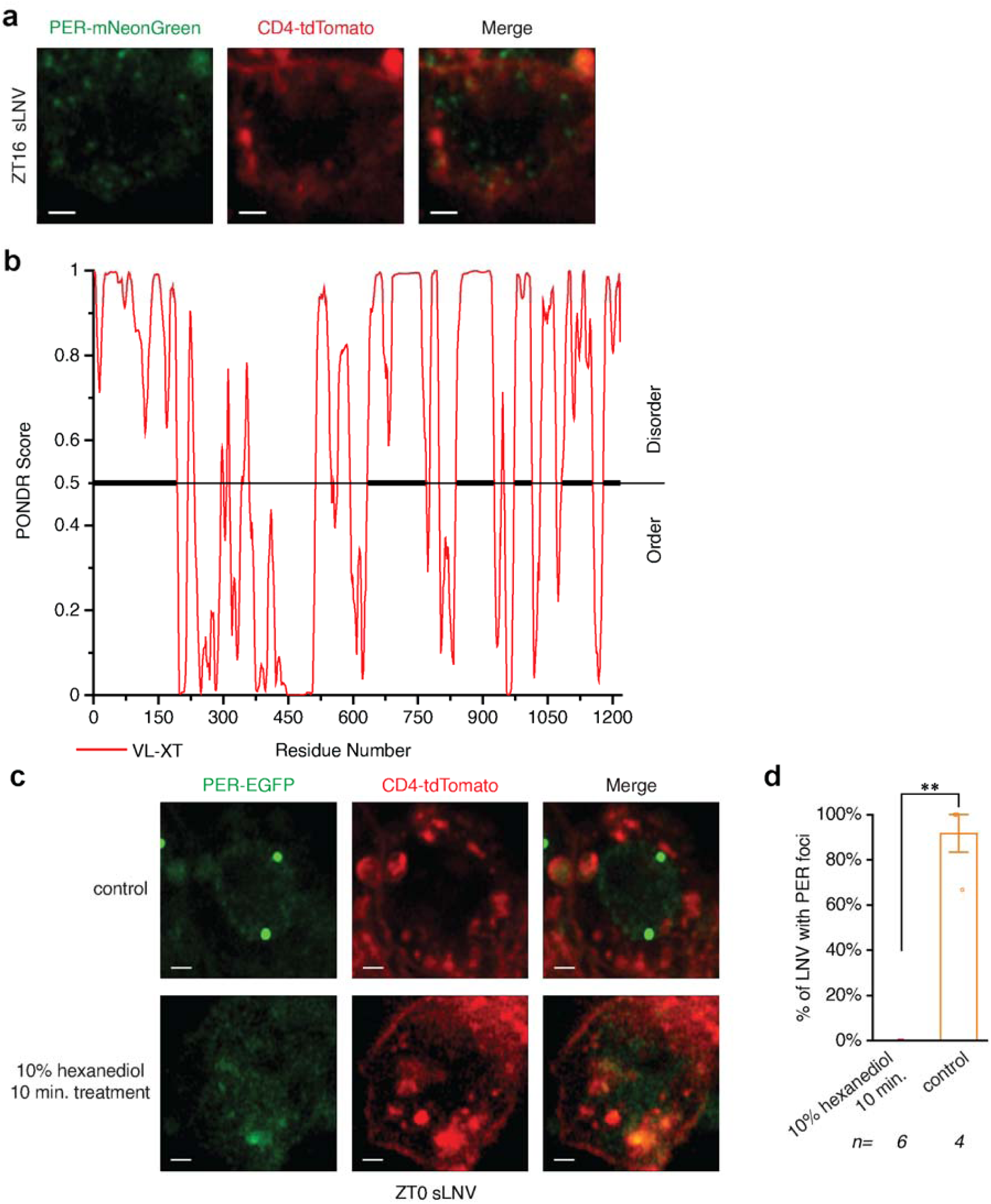
PER foci display liquid-like properties. (a) Representative images of PER foci in the cytoplasm of sLNVs at ZT16 in live imaging experiments. PER foci (green) are present in the cytoplasmic region denoted by CD4-tdTomato (red) staining. (**b**) Analysis of PER amino acid protein sequence. Red line, predictor of natural disordered regions (PONDR) score for intrinsic disorder, >0.5 is considered disordered. (**c**) Representative images of PER foci in sLNvs at ZT0 in control brains and brains treated with 10% 1,6-Hexanediol for 10 minutes. PER foci are disassembled upon hexanediol treatment. (**d**) Quantitation of percentage of sLNvs with PER foci at ZT0 in control and hexanediol treated brains. Scale bars, 1 μm. Statistical test used was a Mann-Whitney test (d). \**P* < 0.05, \*\**P* < 0.005, \*\*\**P* < 0.0005, \*\*\*\**P* < 0.0001. Individual data points, mean, and s.e.m. are shown. See Supplementary Table 2 for detailed statistical analysis.

**Extended Data Fig. 5:**
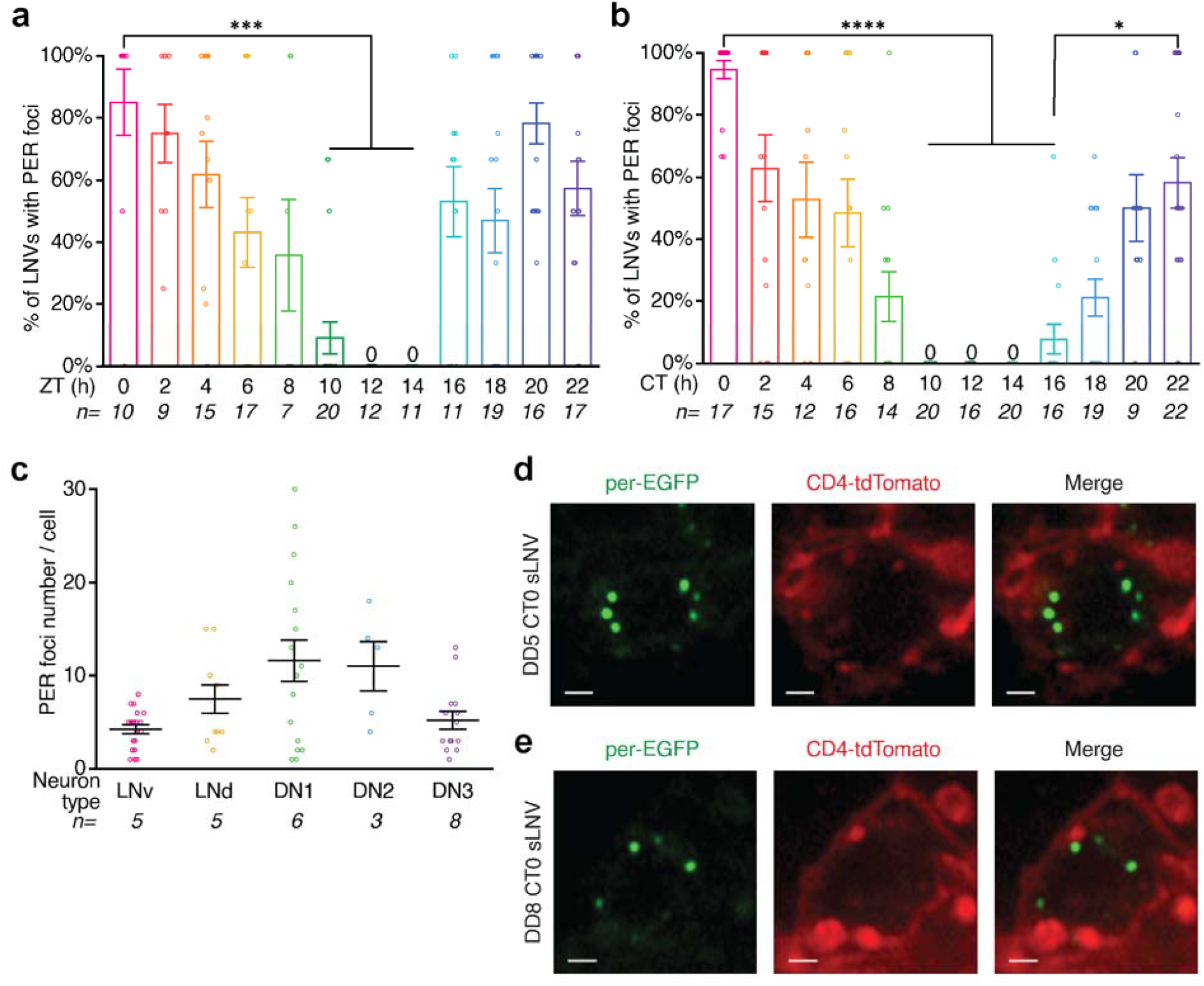
Quantification of PER foci number during the repression phase. (**a** and **b**) Percentage of LNvs with PER foci at different ZTs (a) and CTs (b) over the circadian cycle. ZT denotes time in LD cycles, CT denotes time in constant conditions. PER foci are present in a majority of the LNvs during peak repression phase. (**c**) Quantitation of foci number per different classes of clock neurons at ZT0. (**d** and **e**) *per-EGFP;clk>CD4-tdTomato* flies are entrained to Light-Dark cycles (LD) for 5 days and released into complete darkness (DD). Representative images of sLNVs with PER foci at CT0 on day 5 (d) and day 8 (e) of constant conditions in complete darkness after entrainment to LD cycles. CT refers to ‘Circadian Time’. PER foci persist during the repression phase for several days in constant conditions (complete darkness). Scale bars, 1 μm. Statistical test used were Kruskal-Wallis tests (a, b, and c). \**P* < 0.05, \*\**P* < 0.005, \*\*\**P* < 0.0005, \*\*\*\**P* < 0.0001. Individual data points, mean, and s.e.m. are shown. See Supplementary Table 2 for detailed statistical analysis.

**Extended Data Fig. 6:**
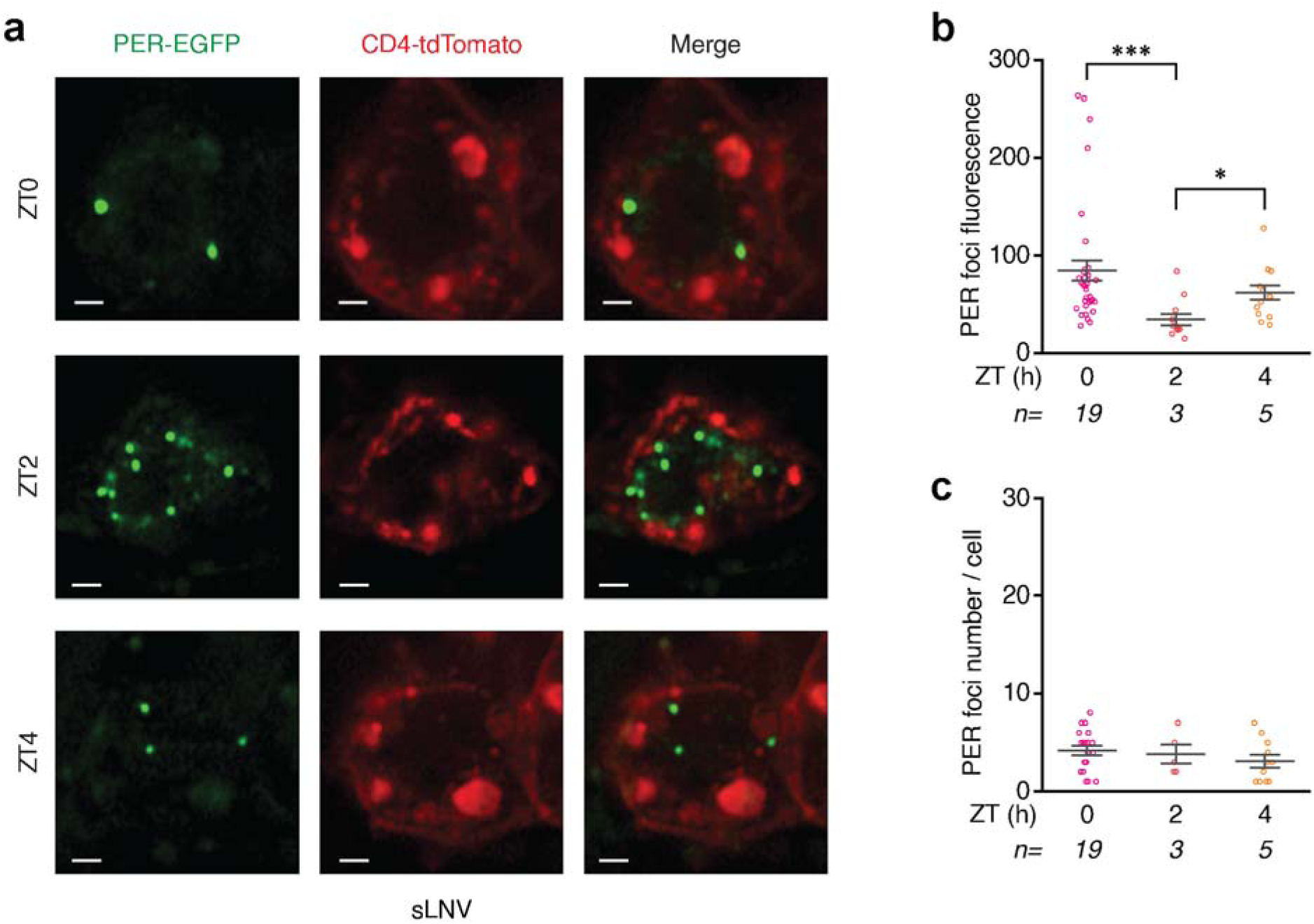
Characterization of PER-EGFP foci in clock neurons. (**a**) *per- EGFP;clk>CD4-tdTomato* flies are entrained to Light-Dark cycles (ZT0-lights on, ZT12-lights off) and live imaging of brains were performed at different times over the circadian repression phase. Shown here are representative images of PER (green) in a class of clock neurons, sLNVs (cell membrane labeled with tdTomato and shown in red) at different ZTs. (b and c) Quantitation of PER-EGFP fluorescence (b) and foci number per sLNV (c) at specific ZTs over the circadian repression phase. Statistical tests used were a Kruskal-Wallis test (**b** and **c**). \**P* < 0.05, \*\**P* < 0.005, \*\*\**P* < 0.0005, \*\*\*\**P* < 0.0001. Individual data points, mean, and s.e.m. are shown. See Supplementary Table 2 for detailed statistical analysis.

**Extended Data Fig. 7:**
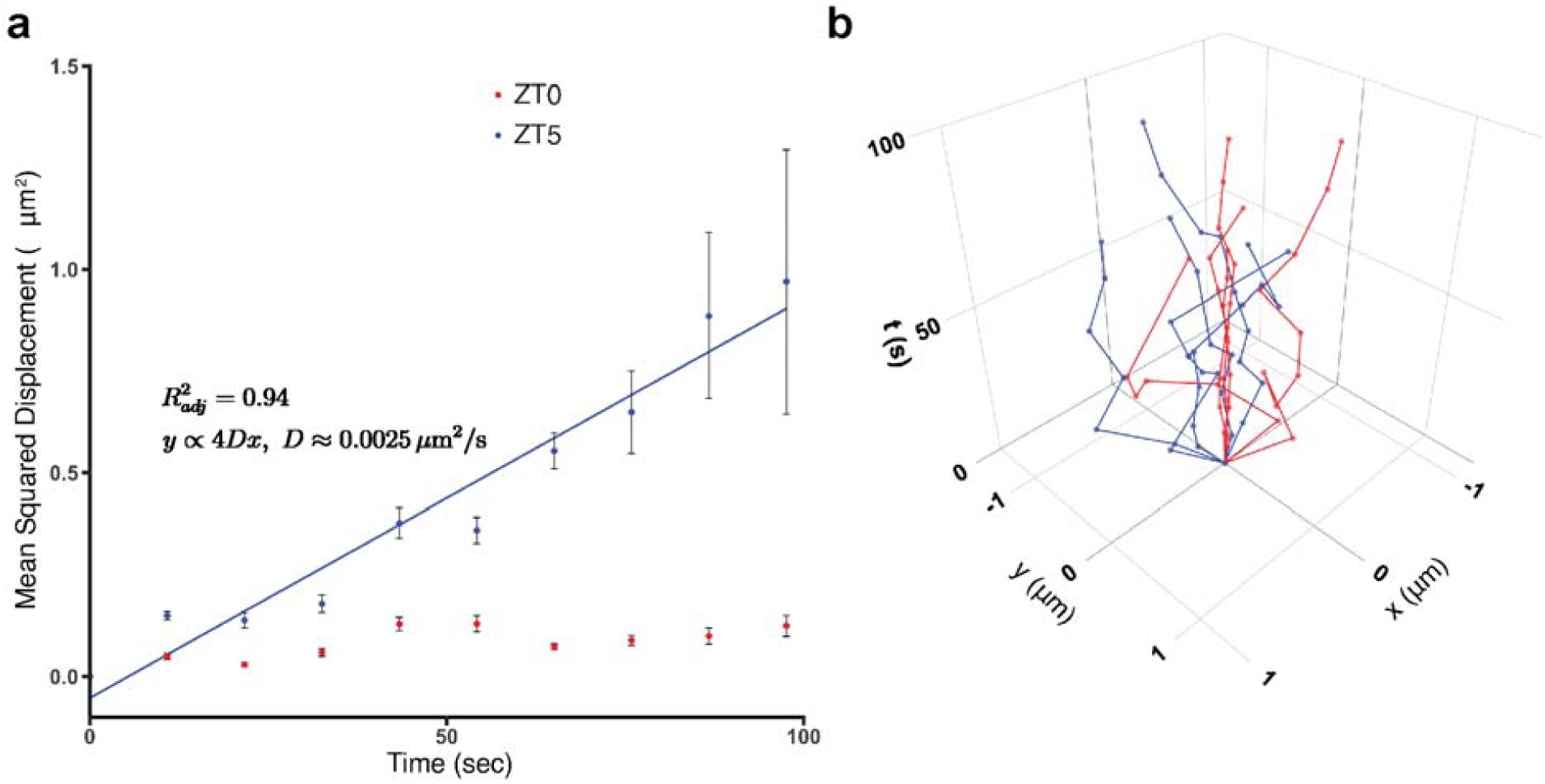
PER foci at ZT0 are more constrained in movement compared to ZT5. (**a**) Mean squared displacement (MSD) of PER foci at different times (ZT0-red dots, ZT5- blue dots) during the repression phase. MSD at ZT5 is fitted using linear regression, suggesting that PER foci at ZT5 are diffusing without a preferred direction (D ≈ 2.5 x 10^-3^ μm^2^/s) . At ZT0, MSD values are significantly lower compared to ZT5 and could not be fitted with a linear model (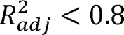), suggesting that PER foci are more constrained at ZT0. Shown here are 0.8 mean and s.e.m. of MSD of foci at ZT0 and ZT5. (**b**) Representative tracks of PER foci movement at ZT0 and ZT5 (ZT0-red, ZT5-blue). X and Y coordinates of individual foci are plotted as a function of time (Z-axis) for each group. See Supplementary Table 2 for details of foci count, mean and s.e.m. values.

**Extended Data Fig. 8:**
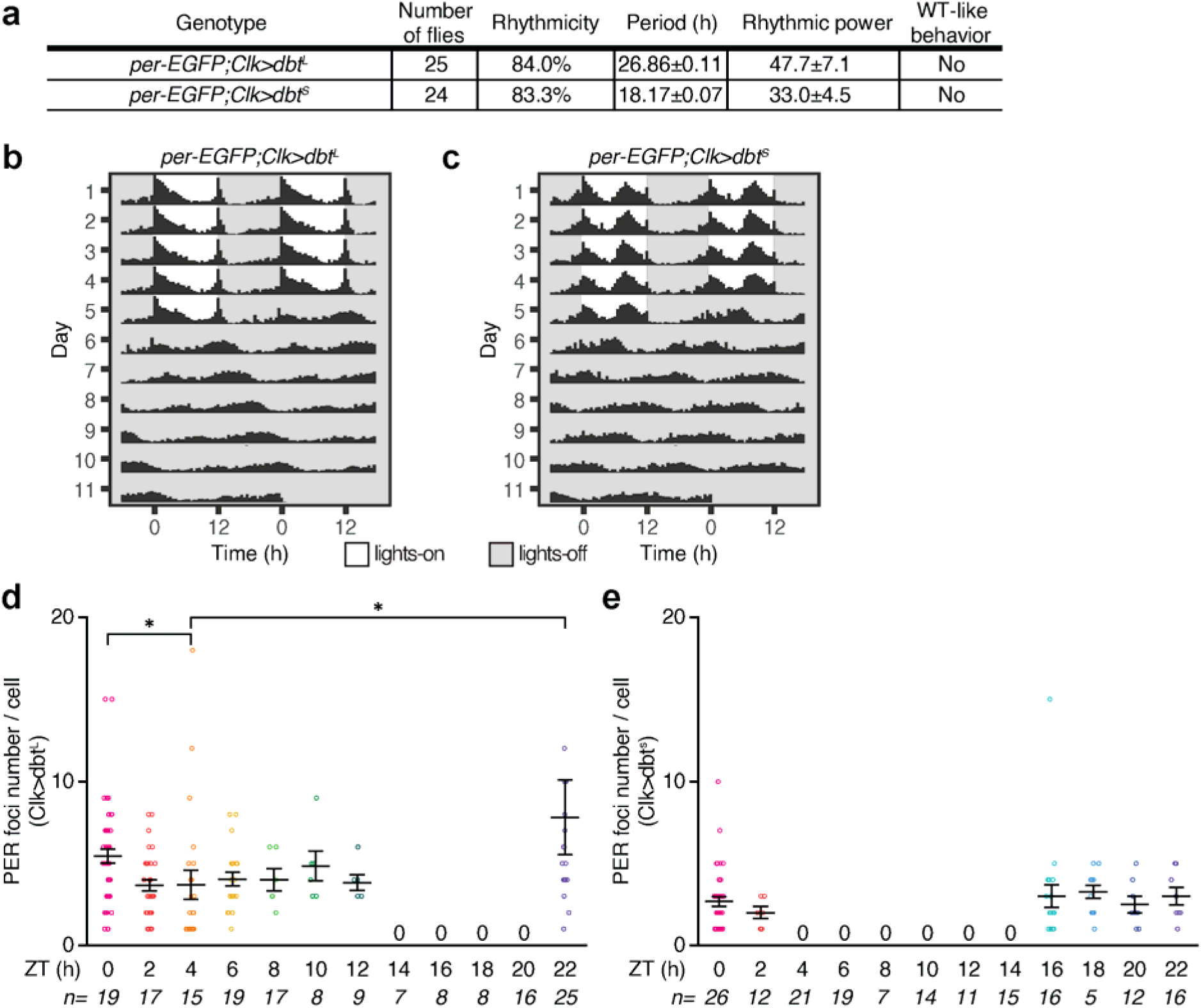
Characterization of PER period foci in *dbt* mutants. (**a**) Summary of free-running locomotor activity rhythms under constant conditions after entrainment to Light- Dark (LD) cycles. The genotypes of the flies tested are listed in the table. Period was calculated by chi-square periodogram analysis for all the flies and shown as average period ± s.e.m.. (**b** and **c**) Flies were entrained to LD cycles (ZT0-lights on, ZT12-lights off) for 5 days and released into constant darkness for 7 days. Averaged population locomotor activity profiles of flies with rest- activity shown for two consecutive days in the same line. *per-EGFP;Clk>dbt^L^* flies display ∼27 hour behavioral rhythms and *per-EGFP;Clk>dbt^S^* flies display ∼18 hour behavioral rhythms. (**d** and **e**) Quantitation of PER foci number in LNv clock neurons at different ZTs over the circadian cycle in *per-EGFP;Clk>dbt^L^* (d) and *per-EGFP;Clk>dbt^S^* (e) flies. Scale bars, 1 μm. Statistical tests used were a Kruskal-Wallis test (d and e). \**P* < 0.05, \*\**P* < 0.005. Individual data points, mean, and s.e.m. are shown. See Supplementary Table 2 for detailed statistical analysis.

**Extended Data Fig. 9:**
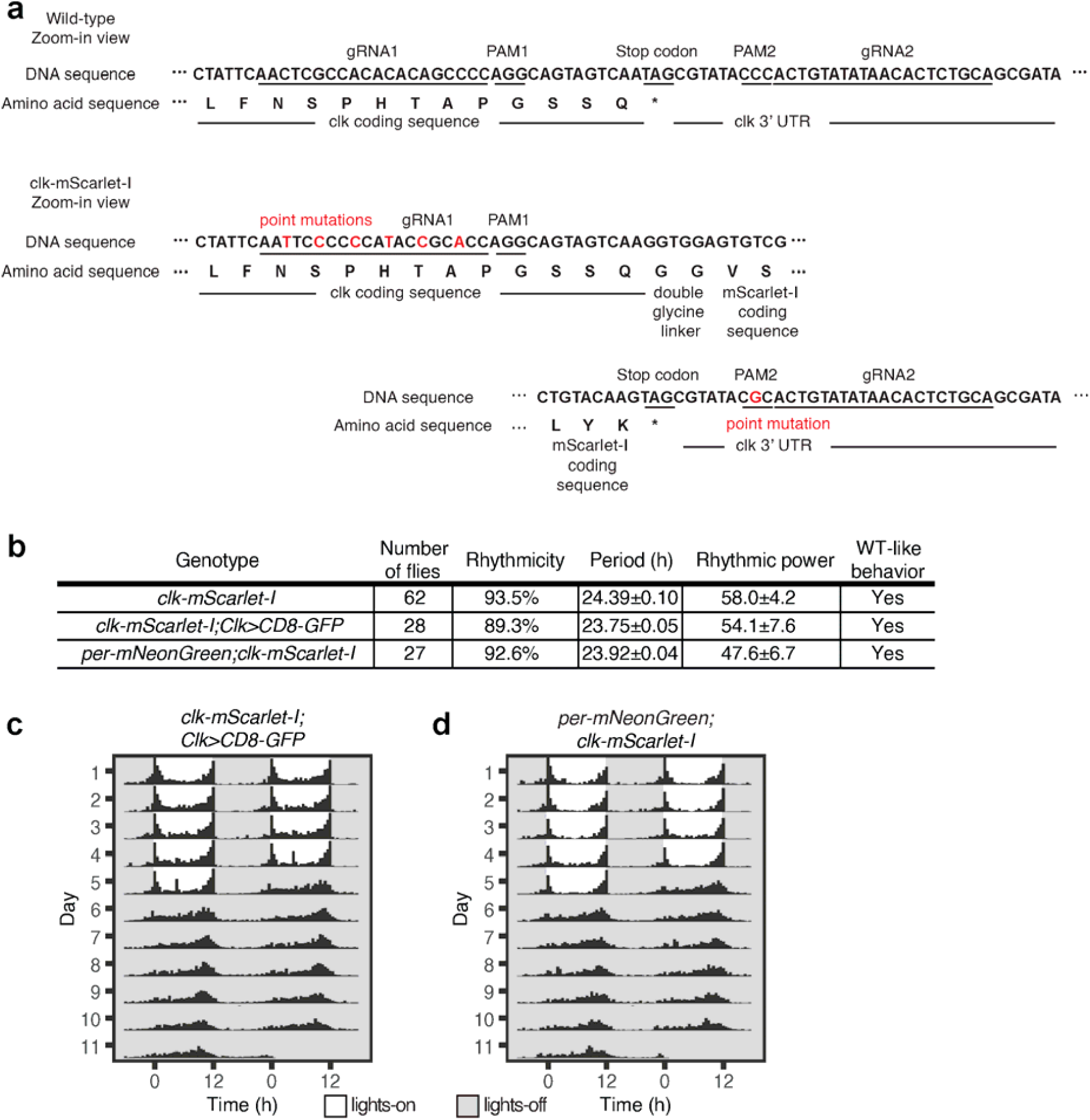
Characterization and behavioral rhythms of *clk-mScarlet-I* flies. (**a**) Shown are the sequencing results of *clk*-mScarlet-I allele. The protospacer adjacent motifs (PAM) of gRNA2 in 3’UTR was mutated (amino acid sequence of the protein was not changed) to prevent cutting by Cas9 after homology directed repair events. PAM sequence of gRNA1 in the last exon was not amenable to mutation, as it would result in change of the amino acid sequence of the protein. Therefore, we mutated multiple other bases in the gRNA1 sequence in order to prevent cutting by Cas9 after homology directed repair. Point mutations are marked in red. (**b**) Summary of free-running locomotor activity rhythms of *clk-mScarlet-I* flies under constant conditions after entrainment to Light-Dark (LD) cycles. Period was calculated by chi- square periodogram analysis for all the flies and shown as average period ± s.e.m.. (**c**, and **d**) Flies of indicated genotypes were entrained to LD cycles (ZT0-lights on, ZT12-lights off) for 5 days and released into constant darkness for 7 days. Averaged population locomotor activity profiles of flies with rest-activity shown for two consecutive days in the same line. These flies display rhythmic behaviors with activity peaks around the time of lights-on and lights-off.

**Extended Data Fig. 10:**
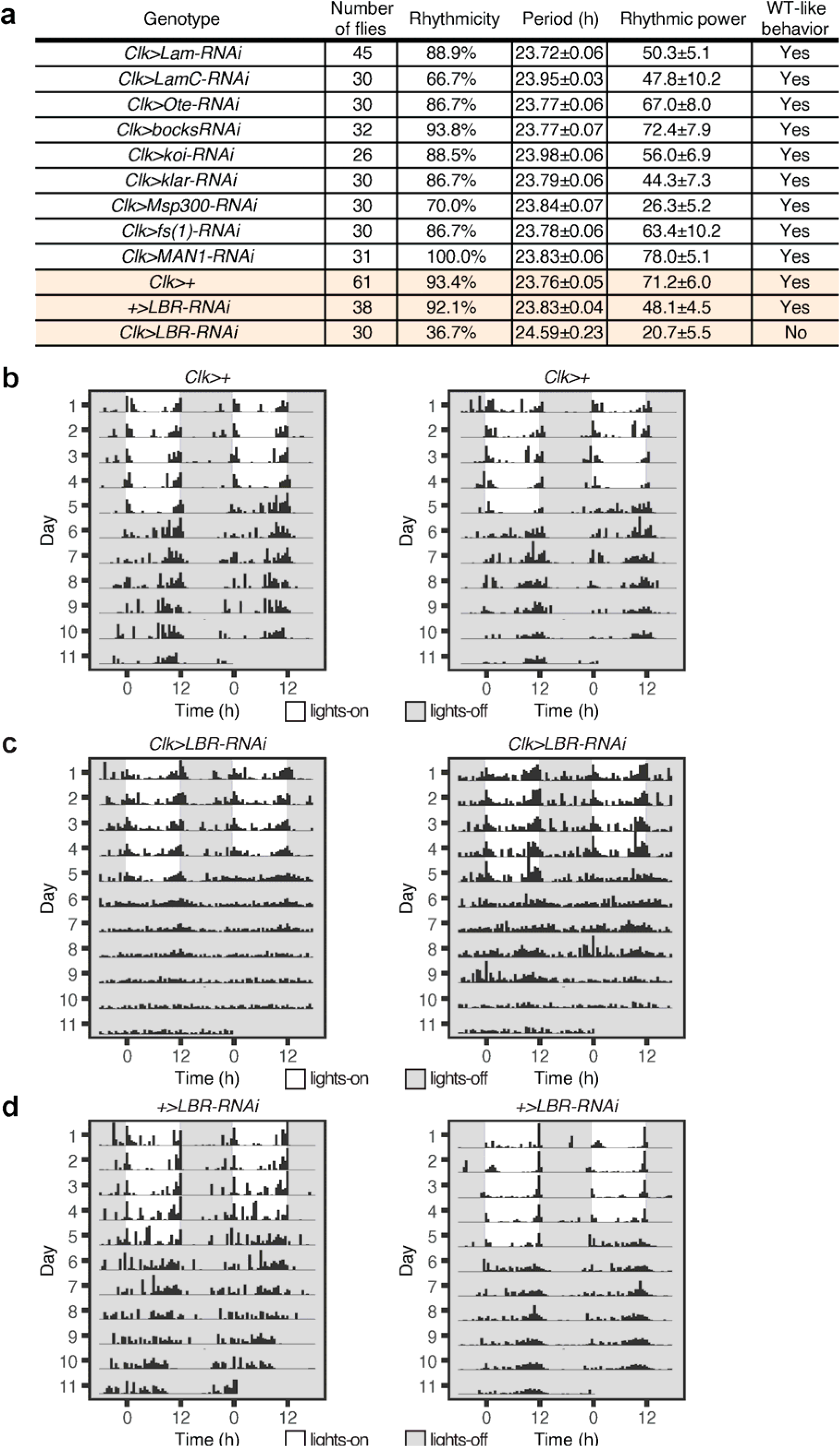
Summary of locomotor activity rhythms and periodicity in flies in which LBR is knocked down in clock neurons. (**a**) Summary of free-running locomotor activity rhythms of *Clk-Gal4>UAS-RNAi* flies under constant conditions after entrainment to Light-Dark (LD) cycles. Period was calculated by chi-square periodogram analysis for all the flies and shown as average period ± s.e.m.. (**b, c** and **d**) Lamin B receptor (LBR) expression is knocked down specifically in the clock neurons by crossing *Clk-Gal4* flies to *LBR-RNAi* flies. Control flies (*Clk-Gal4>+, +>UAS-LBR-RNAi*) and experimental flies (*Clk-Gal4>UAS-LBR- RNAi*) were entrained to LD cycles (ZT0-lights on, ZT12-lights off) for 5 days and then released into complete darkness for 7 days. Representative activity profile of individual *Clk-Gal4>+* flies (b), *Clk-Gal4>UAS-LBR-RNAi* (c), and *+>UAS-LBR-RNAi* flies (d) with rest-activity shown for two consecutive days in the same line. Control flies show rhythmic behavior in constant conditions (complete darkness) with distinct morning and evening activity peaks, while *Clk- Gal4>UAS-LBR-RNAi* flies display arrhythmic behavior with no distinct activity peaks.

## Materials and Methods

### Data reporting

We followed standard protocols to conduct our *Drosophila* live imaging and circadian behavior experiments. No statistical calculations were used to pre-determine sample sizes. Our sample sizes are similar to those generally employed in the field. Samples were not randomized for the experiments and blinding was not used. We reported the sample sizes for each of our experiments in the Supplementary table files.

### Fly strains

Flies were raised on cornmeal-yeast-sucrose food under a 12:12 Light:Dark cycle at 25 °C and 60-70% humidity. The following flies used in the study were previously described or obtained from the Bloomington Stock Center: *w^1118^* (BL-6326), *y,sc,v* (BL-25709), *Clk-Gal4*(29) (expressed in all clock neuron classes), *elav-Gal4*(63), *per^01^* (28), *UAS-CD8GFP*(30), *UAS- CD4tdTom*(30), *UAS-dbt^L^* (38), *UAS-dbt^S^* (38), *UAS-Lam-RNAi* (BL-57501), *UAS-LamC-RNAi* (BL-31621), *UAS-bocks-RNAi* (BL-38349), *UAS-Ote-RNAi* (BL-39009), *UAS-fs(1)Ya-RNAi* (BL-64597), *UAS-MAN1-RNAi* (BL-65371), *UAS-koi-RNAi* (BL-40924), *UAS-klar-RNAi* (BL-36721), *UAS-Msp300-RNAi* (BL-32848), *UAS-LBR-RNAi* (BL-53269). The following flies were obtained as indicated: *UAS-unc84-tdTomato*(36) (H. Gilbert, HHMI/Janelia Farm Research Campus), *period*-AID-EGFP - referred to in the paper as *per-EGFP*(35) (Y. Zhang, University of Nevada, Reno). Specific genotypes of flies used in the experiments are presented in the figures and in the Supplementary Table 2 file. For control experiments, Gal4 and UAS lines are crossed to *w^1118^* flies (the genetic background of Gal4 and RNAi lines).

### Protein disorder analysis

*Drosophila* PER protein amino acid was obtained from flybase. Protein disorder was predicted using the program Predictor of Natural Disorder Regions (PONDR) version VL-XT(64).

### CRISPR protocol for generating *per-mNeonGreen* and *clk-mScarlet-I* flies

The protocol of sgRNA plasmid generation for per-mNeonGreen and clk-mScarlet-I knock-in is based on U6-gRNA (chiRNA) cloning protocols (https://flycrispr.org/protocols/gRNA/). For each fly line, two sgRNAs are used for higher efficiency. Positive sgRNA plasmids were confirmed by Sanger sequencing and prepared for injection (QIAGEN Plasmid Midi Kit). Rainbow Transgenic Flies, Inc. performed all injections into *nos-Cas9* attp2 fly embryos. Supplementary Table 1 provides details of synthesized long fragments, cloning primers, single-guide RNA (sgRNA) sequences, and sequencing primers.

#### per-mNeonGreen fly

To generate *per-mNeonGreen* fly line, the donor plasmid was constructed via Golden Gate Assembly. First, the 5’-homology arm was amplified from *y,sc,v* genomic DNA using Q5 High-Fidelity DNA Polymerase (New England BioLabs, NEB) with primers: per-5’- pcrF and per-5’-pcrR (listed in Supplementary Table 1), and then purified (QIAGEN, PCR purification kit). The 3’-homology arm (∼1 kb) was directly synthesized by IDTDNA. The mNeonGreen sequence (based on Addgene Plasmid #98886) was codon-optimized (IDT, Codon Optimization Tool) and directly synthesized from IDTDNA. Subsequently, EcoRI, XbaI-linearized pBluescript sk(-) plasmid backbone was mixed with 5’ arm, mNeonGreen fragment, and 3’ arm fragments. Golden Gate Assembly protocol was applied with type IIS endonuclease Esp3I (NEB). The product was transformed into competent E. coli cells (NEB, C2987H) and plated on LB-Amp plates. Single colonies were screened by colony PCR using DreamTaq Green PCR Master Mix (Thermo Scientific, K1082) with primers: seq-per-F and seq-per-R (listed in Supplementary Table 1). Positive colonies were purified (QIAGEN, Plasmid Mini Prep Kit), confirmed by Sanger sequencing and prepared for injection (QIAGEN, Plasmid Midi Kit).

After injected embryos developed into adult flies, these adult flies were individually crossed to FM7a male or virgin female flies, as *period* gene is on X chromosome and FM7a is a X chromosome balancer. We PCR screened 126 progenies (F1) for the presence of the ∼700bp insert, and 47 of them were positive. We followed the protocol described here for screening: Genomic DNA was extracted from the F1 flies by removing a midleg and placing it into the well of a 96-well plate containing 10 µl of lysis buffer (10 mM Tris pH 8.2, 1 mM EDTA pH 8.0, 25 mM NaCl, 400 µg ml^−1^ Proteinase K). Each fly was then placed in the corresponding well of a 96-well deep-well plate (Brandtech VWR, 80087-070) filled halfway with fly food and capped with cotton. The 96-well plate containing lysis buffer and fly legs was then incubated at 37 °C for 1.5 hours followed by a 5-min heat inactivation step at 95 °C. Genomic DNA (1.5 µl) from the reaction was used as the PCR template for a 10-µl reaction of DreamTaq Green PCR Master Mix (Thermo Scientific, K1082) for 35 cycles. The product from the PCR reaction was run on a 1% agarose gel. We backcrossed the flies that had the ∼700bp insertion to FM7a flies and then homozygosed their progeny. Homozygous stocks were genotyped and Sanger-sequenced.

#### Clk-mScarlet-I fly

To generate *clk-mScarlet-I* fly line, the donor plasmid was constructed via Golden Gate Assembly steps. 3’ homozygous arm was amplified from *y,sc,v* genomic DNA using Q5 High-Fidelity DNA Polymerase (New England BioLabs, NEB) with primers: clk-3’- pcrF and clk-3’-pcrR (listed in Supplementary Table 1), and then purified (QIAGEN, PCR purification kit). 5’ homologous arm was amplified through overlap extension: Fragment 1 was amplified from *y,sc,v* genomic DNA using Q5 High-Fidelity DNA Polymerase (New England BioLabs) with primers- clk-5’-Fragment1-F and clk-5’-Fragment1-R (listed in Supplementary Table 1). Fragment 2 was directly synthesized from IDT. Fragments 1 and 2 were mixed and amplified with primers: clk-5’-Fragment1-F and clk-5’-Fragment2-R (listed in Supplementary Table 1). Then, the overlap extension product was purified (QIAGEN, PCR purification kit). The mScarlet-I sequence (based on Addgene Plasmid #85068) was codon-optimized (IDT, Codon Optimization Tool) and directly synthesized from IDTDNA. Golden Gate Assembly steps were performed as described above to generate clk-mScarlet-I donor plasmid, which was then prepared for injection.

The adult injected flies were individually crossed to TM6/TM3 male or virgin female flies as *clock* gene is on 3^rd^ chromosome and TM6, and TM3 are 3^rd^ chromosome balancers. We PCR screened 149 progenies (F1) for the presence of the ∼700bp insert, and 34 of them were positive. We performed similar steps as described above to screen for *clk-mScarlet-I* flies and generated homozygous stocks.

### Live imaging protocol

Flies were entrained to Light-Dark (LD) cycles with lights on for 12 hours and off for 12 hours for 5-7 days, and then released into complete darkness (DD) for 6-7 more days. Zeitgeber Time (ZT) 0 marks the beginning of the light phase, and ZT12 marks the onset of the dark phase. Circadian time (CT) refers to times in complete darkness (DD) and CT0 is the start of the subjective light phase and CT12 is the start of the subjective dark phase, the times when the light transitions would have occurred had the LD cycle continued. For instance, DD1 is the first day in complete darkness and DD2 is the second day in incomplete darkness.

All flies used for live imaging experiments were placed in density-controlled food vials (4 females and 4 males) and entrained for 5-7 days in incubators. We performed all our live imaging experiments on 5-7 day old male or female flies, and did not notice any differences in our experimental results. The fly genotype and ZT was as described in the figure legends. We used the GAL4/UAS system to express transgenes in clock neurons in the brain. We imaged our samples using a Zeiss LSM800 laser scanning confocal microscope with AiryScan super-resolution module (125 nm lateral and 350 nm axial resolution). We have acquired our images using a 63x Plan-Apochromat Oil (N.A. 1.4) objective and 405, 488, and 561 nm laser lines. We collected Z-stack (each Z-slice is about 250 nm) or time-lapse image series of individual clock neurons and images were analyzed using Zeiss ZEN software and ImageJ.

#### per-mNeonGreen and per-EGFP imaging

We generated flies with a red fluorescent cell membrane marker throughout the circadian clock network by crossing *Clk-Gal4* (pan-clock neuron driver) to *UAS-CD4-tdTomato* (red membrane marker) flies. To visualize the inner nuclear membrane, we expressed *UAS-unc84-tdTomato* in all clock neurons by crossing it with the *Clk-Gal4* driver. unc84 is a *C. elegans* SUN domain protein and *unc84-tdTomato* localizes red fluorescent protein to the inner nuclear membrane. We then crossed these flies with either *per-mNeongreen* or *per-EGFP* flies to enable PER protein imaging in all the clock neurons in the brain. For *dbt^L^* and *dbt^S^* experiments, we first generated *per-EGFP;Clk-Gal4>UAS-CD4-tdTomato* flies and then crossed them to *UAS-dbt^L^* or *UAS-dbt^S^* flies. For live imaging of clock neurons, 3-4 brains were dissected in chilled Schneider’s *Drosophila* medium (ThermoFisher Scientific, 21720001) in less than 5 minutes. A punched double-sided tape was used as a spacer on the slides to prevent flattening of the brains. The brains were overlaid with a small amount of Prolong^TM^ Glass Antifade mounting medium (ThermoFisher Scientific, P36982) and covered using a coverslip. We acquired individual z-stack images of clock neurons using the Zeiss LSM800 Airyscan laser scanning confocal microscope.

#### clk-mScarlet-I imaging

We generated flies with a green fluorescent cell membrane marker throughout the circadian clock network by crossing *Clk-Gal4* to *UAS-CD8-GFP* (green membrane marker) flies. We then crossed these flies to *clk-mScarlet-1* flies to enable CLK protein imaging in all the clock neurons in the brain. For *per^01^* experiments, we first generated *clk-mScarlet-I;Clk-GAL4>UAS-CD8-GFP;* flies and then crossed them with female *per^01^* flies and we used male flies for our experiments: experimental flies - *per^01^/Y; Clk-Gal4>UAS-CD8- GFP;clk-mScarlet-I*, control flies: *+/Y; Clk-Gal4>UAS-CD8-GFP;clk-mScarlet-I*. We followed similar experimental protocols for entrainment and live imaging as described above.

### Quantitative real-time PCR

At indicated time points, ∼30 flies were collected for each group. Total RNA was extracted from fly heads using Trizol Reagent (Invitrogen, 15596026), and cDNAs were synthesized using TaqMan Reverse Transcription Reagents (Invitrogen, 4304134) using primers: per-R and rp49-R (listed in Supplementary Table 1). Quantitative real-time PCR was performed on QuantStudio3 (Applied Biosystems) using default thermal cycling conditions. Power SYBR Green Master Mix (Applied Biosystems, 4368577) was used with *per* and *rp49* primers listed in Supplementary Table 1. Data were analyzed using the 2^−ΔΔCT^ method with mRNA levels normalized to the gene *rp49*. Relative mRNA amplitude was calculated with respect to the trough (CT3) levels set as 1 for each fly line.

### DNA FISH probe synthesis

We used FISH Tag DNA Green Kit (Invitrogen, F32947) to generate *period* DNA probes and FISH Tag DNA Orange Kit (Invitrogen, F32948) to generate *timeless* DNA probes. In the Drosophila genome, *period* gene is on X chromosome and *timeless* gene is on II^nd^ chromosome. Each set of probes covered 12 to 25 kb genomic region covering the target gene and the adjacent regions. Within each region, multiple ∼1kb or ∼1.5kb fragments were selected based on the following criteria: 1) 39% to 41% total GC content, and 2) existence of primer pairs with similar melting temperatures (±5 °C). Primers used to generate DNA FISH probes are listed in Supplementary Table 1. Next, fragments were amplified from *y,sc,v* genomic DNA using Q5 High-Fidelity DNA Polymerase (New England BioLabs, NEB) with all primers listed in Supplementary Table 1 and then purified (QIAGEN, PCR purification kit). Before nick translation, DNaseI concentration was tested and adjusted to ensure appropriate probe length (∼300 bp). Finally, later steps including nick translation, labeling with fluorescent dye, and purification are performed according to the instructions in the kit manual (Invitrogen, F32947). DNA FISH experiments were conducted using fresh probes to ensure sensitivity and accuracy.

### Immunofluorescent staining in brains

*Drosophila* brains were dissected in chilled Schneider’s *Drosophila* medium. We pooled all the brains that were dissected in 10 minutes for each of our experiments. Brains were fixed for 20 minutes with gentle rocking in 4% formaldehyde in PBS at room temperature. Fixed brains were briefly rinsed three times and permeabilized in PBST (1X PBS + 0.3% Triton X-100) at room temperature for 1 hour before adding primary antibody. Brains were incubated with primary antibody in 5% normal goat serum (NGS) in PBST at 4 °C overnight or for at least 16 hours. Samples were then washed three times for 20 minutes each in PBST. Samples were then incubated with secondary antibody in 5% NGS in PBST overnight at 4 °C or at room temperature for 3-5 hours, and then washed three times for 20 minutes each in PBST. Samples were mounted in Prolong^TM^ Glass Antifade mounting medium (ThermoFisher Scientific, P36982). The following primary antibodies were used (dilutions noted in parentheses): mouse anti-PDF C7(65) (1:500) (Developmental Studies Hybridoma Bank, Iowa), and rabbit anti-PER(66) (1:500, gift from Dr. Orie Shafer). Alexa 488 and Alexa 555 (Molecular probes) fluorescence conjugated secondary antibodies were used at a 1:500 dilution. Images were taken using a Zeiss LSM 800 laser scanning confocal microscope with Airyscan super-resolution detector with a 63x oil immersion objective and processed using Image J.

### Immuno-DNA FISH protocol

Brains were first subjected to immunostaining steps and then DNA-FISH steps. For the immunostaining step, we followed the same protocol as described above. DNA-FISH protocol is adapted from a previous study(62). Briefly, brains were dissected in chilled Schneider’s *Drosophila* medium and fixed in 4% paraformaldehyde in 1X PBS for 20 minutes. Samples were then washed, and permeabilized for at least 1 hour in PBST, and incubated with primary antibodies overnight at 4°C. Samples were then washed with PBST (20 min, three times), incubated with Alexa Fluor-conjugated secondary antibodies (1:200; Molecular Probes) overnight at 4°C, and washed again with PBST (20 min, three times). Samples are protected from light for all the next steps starting from the secondary antibody addition step. Samples were then fixed for 10 min with 4% formaldehyde in 1X PBS followed by three washes in PBST for 5 min each. Samples were then treated with RNase A (2 mg/ml in water, Qiagen, 19101) for 2 hours at 37°C, washed with PBST for 15 min, and stained with 1 μg/mL of Hoechst 33258 (Sigma Aldrich, 94403) for 15 min at room temperature. To allow gradual transition into 50% formamide solution (50% formamide, 4X SSC, 100mM NaH_2_PO_4_, 0.1% Tween20) samples were incubated sequentially for a minimum of 10 min each in 20% formamide solution (20% formamide, 4X SSC, 100mM NaH_2_PO_4_, 0.1% Tween20), 40% formamide solution (40% formamide, 4X SSC, 100mM NaH_2_PO_4_, 0.1% Tween20), and finally in 50% formamide solution at room temperature. The brains were then incubated in 50% formamide solution at 80°C for 15 minutes. During the final step, the fluorescence-labeled DNA probes (70-100 ng/μl) were added to the hybridization mixture (50% formamide, 2X SSC, 10% dextran sulfate, 0.5 mg/ml Salmon sperm DNA) and the solution was incubated at 95 °C for 3 minutes prior to hybridization. The hybridization solution was added to the samples and hybridization was carried out at 37°C overnight. Following hybridization, samples were washed twice in 50% formamide/2X SSCT for 30 minutes each at 37 °C, once in 40% formamide/2X SSCT and once in 20% formamide/2X SSCT and three times in 2X SSCT with each wash for 10 minutes all at room temperature. Samples were mounted using Prolong^TM^ Glass Antifade mounting medium and images were acquired using a Zeiss LSM800 confocal microscope with a 63x oil immersion objective.

### Locomotor activity and Rhythmicity analysis

Individual adult male flies (3-5 days old) were placed in glass capillary tubes (∼4 mm inner diameter, 5 cm in length) containing 2% agar and 4% sucrose food, which were then loaded into TriKinetics DAM2 *Drosophila* Activity Monitors (Waltham, MA, USA) for locomotor activity recordings. Flies were entrained to Light-Dark (LD) cycles with lights on for 12 hours and off for 12 hours for 5-7 days, followed by complete darkness (DD) for six more days. The DAM monitors are equipped with infrared sensors and they record infrared beam breaks when the flies cross the middle of the glass tube. The monitors with flies were placed in the incubators and the beam crossing counts from each monitor were recorded on a computer. The Beam crossing counts were placed into 30-minute bins for time-series analysis of locomotor activity. Averaged population activity profiles under Light-Dark cycles and constant conditions were generated using a commercially available software ClockLab (Actimetrics) and public domain R Rethomics software package. Briefly, activity levels were normalized for each fly, such that the average number of beam crossings in each day (48 bins) is equal to 1. Next, the population average of normalized activity was determined and the results are displayed as normalized activity plots in the figures. We used activity counts of individual flies under complete darkness conditions following LD cycles to analyze rhythmicity and to determine the free-running period of the circadian clock. Rhythmicity and free-running period of individual flies were determined by a chi-square periodogram analysis with a confidence level of 0.001 using the ClockLab software. The “Power” and “Significance” values generated from the chi-square analysis were used to calculate “Rhythmic Power” as a measure of the strength of each rhythm.

### 1,6-Hexanediol Treatment of *Drosophila* brains

For treatment with 1,6-Hexanediol, Schneider’s *Drosophila* medium was prepared containing 10% 1,6-Hexanediol (Sigma-Aldrich, 240117). The brains were dissected in Schneider’s *Drosophila* medium and then incubated in 10% 1,6-Hexanediol medium or control medium (Schneider’s) for 10 minutes at room temperature. The brains were then mounted on a slide and imaged immediately following the treatment.

### Data analysis

A custom ImageJ (Fiji distribution) Python plugin was developed in order to automate image data analysis. In our live imaging experiments, Z-stack or time-lapse series were captured by Zeiss ZEN software and subjected to channel alignment, if necessary. CZI files were then imported into ImageJ using Bio-Formats Importer as a composite image with a lossless 16-bit resolution per channel. Next, regions of interest (ROIs) are manually annotated on the channel using appropriate white values for visualization. We used the same visualization settings for all images from the same fly line and neuron type to ensure unbiased fluorescence analysis.

#### Analysis of foci fluorescence

First, we selected a single Z-plane with the largest foci count for each cell. Next, each individual foci was annotated with an elliptical ROI and its fluorescence value was calculated by subtracting the background fluorescence from the integrated intensity of the ROI region (ImageJ). Background fluorescence per pixel was estimated by the mean intensity of a background selection on the same plane.

#### Analysis of distance between PER protein foci or period gene and the nuclear periphery

First, we selected the center Z-plane for each cell where the nuclear envelope or DAPI staining is clearly visible. Next, foci were annotated with elliptical ROIs. In the same center Z-plane, nuclear periphery was annotated with closed polygon ROIs, based on DAPI/unc84-tdTomato signals. Finally, we calculated the shortest distance from the centroid of each foci to the respective closed polygon. We have used similar analysis techniques for calculating the distance between PER protein foci and the nuclear envelope and between *period* gene and the nuclear periphery.

#### Analysis of foci count and percentage of cells with foci

Both analyses were performed independently by manually counting foci across all Z-planes for each cell.

### Movement Analysis

For each cell, an appropriate Z-plane in which PER foci were clearly visible was chosen. A time-lapse image was taken every 10 seconds to capture foci movement. Each individual foci was annotated with an elliptical ROI. For each focus, we tracked its centroid position in each frame and output a time series of centroid coordinates. Next, a custom R script was used to calculate mean squared displacement in the fixed Z-plane. For both ZT time points, we chose foci that were visible for 60-100 seconds. The diffusion coefficient of foci at ZT5 was estimated by least-squares fitting using MSD α 4*Dt*, where D is the diffusion coefficient. Finally, ten foci movement tracks (five from each ZT group) were randomly sampled and plotted using the R Plotly package to show representative traces.

### Statistics and Reproducibility

To analyze foci characteristics (including foci fluorescence, percentage of cells with foci, and foci number per cell), the data is collected from a pool of biological replicates of hemi-brain images collected from more than 3 independent experiments. Fluorescence measurements were taken from distinct neurons from a brain and same neurons were not measured repeatedly. We have used 3 biological replicates for our qPCR and other biochemical experiments with ∼30 flies for each experiment and 30-60 flies for each of our behavior experiments. Data for DNA FISH experiments were collected from more than 3 independent experiments. We have used 3-5 day old male and female flies in all our live imaging and biochemical experiments and only male flies for our behavior experiments. We used OriginPro from OriginLab (Northampton, MA, USA) to graph and statistically analyze data. We used non-parametric tests because the values were not normally distributed as reported by D’Agostion–Pearson omnibus and Shapiro–Wilk normality tests. We adjusted *P* values accordingly when multiple comparisons were conducted. See Supplementary Table 2 for more details on number of experiments, statistical tests, and *P* values for all the experiments.

## Supplemental Tables

**Supplementary Table 1.** Per-mNeonGreen and Clk-mScarlet-I CRISPR information, primers used for Quantitative real-time PCR, and primers used to generate DNA FISH probes in this study.

**Supplementary Table 2.** Source file with details of statistical analysis for experiments related to all the figures.

## Supplemental Movies

**Movie 1.** Time lapse video of sLNVs at ZT0 from *per-mNeonGreen;Clk-Gal4>UAS-CD4- tdTomato* flies (ZT0-lights on). PER foci are shown in green, and clock cell membrane is shown in red.

**Movie 2.** Another representative time lapse video of sLNVs at ZT0 from *per-mNeonGreen;Clk- Gal4>UAS-CD4-tdTomato* flies (ZT0-lights on). PER foci are shown in green, and clock cell membrane is shown in red.

**Movie 3.** Time lapse video of sLNVs at ZT5 from *per-mNeonGreen;Clk-Gal4>UAS-CD4- tdTomato* flies (ZT0-lights on). PER foci are shown in green, and clock cell membrane is shown in red.

**Movie 4.** Time lapse video of PER foci fusion in sLNVs at ZT22 from *per-mNeonGreen;Clk- Gal4>UAS-CD4-tdTomato* flies (ZT0-lights on). PER foci are shown in green, and clock cell membrane is shown in red.

**Movie 5.** Z-stack video of sLNVs at ZT0 from *per-EGFP;Clk-Gal4>UAS-unc84-tdTomato* flies (ZT0-lights on). Video is compiled from individual z-stacks. PER foci are shown in green, and nuclear envelope of sLNVs is shown in red.

**Movie 6.** Another z-stack video of sLNVs at ZT0 from *per-EGFP;Clk-Gal4>UAS-unc84- tdTomato* flies (ZT0-lights on). PER foci are shown in green, and nuclear envelope of sLNVs is shown in red.

**Movie 7.** Z-stack video of sLNVs at ZT5 from *per-EGFP;Clk-Gal4>UAS-unc84-tdTomato* flies (ZT0-lights on). PER foci are shown in green, and nuclear envelope of sLNVs is shown in red.

## Notes

### Competing Interest Statement

The authors have declared no competing interest.

## REFERENCES

1. Takahashi JS (2017) Transcriptional architecture of the mammalian circadian clock. Nat. Rev. Genet. 18(3):164–179.

2. Stanewsky R, et al. (1998) The cryb mutation identifies cryptochrome as a circadian photoreceptor in Drosophila. Cell 95(5):681–692.

3. Buhr ED, Yoo SH, & Takahashi JS (2010) Temperature as a universal resetting cue for mammalian circadian oscillators. Science 330(6002):379–385.

4. Yadlapalli S, et al. (2018) Circadian clock neurons constantly monitor environmental temperature to set sleep timing. Nature 555(7694):98–102.

5. Hardin PE, Hall JC, & Rosbash M (1990) Feedback of the Drosophila period gene product on circadian cycling of its messenger RNA levels. Nature 343(6258):536–540.

6. Aronson BD, Johnson KA, Loros JJ, & Dunlap JC (1994) Negative Feedback Defining a Circadian Clock - Autoregulation of the Clock Gene-Frequency. Science 263(5153):1578–1584.

7. Sehgal A, Price JL, Man B, & Young MW (1994) Loss of circadian behavioral rhythms and per RNA oscillations in the Drosophila mutant timeless. Science 263(5153):1603–1606.

8. Liang XT, Holy TE, & Taghert PH (2016) Synchronous Drosophila circadian pacemakers display nonsynchronous Ca^2+^ rhythms in vivo. Science 351(6276):976–981.

9. Stoleru D, Peng Y, Agosto J, & Rosbash M (2004) Coupled oscillators control morning and evening locomotor behaviour of Drosophila. Nature 431(7010):862–868.

10. Aguilar-Arnal L, et al. (2013) Cycles in spatial and temporal chromosomal organization driven by the circadian clock. Nat. Struct. Mol. Biol. 20(10):1206–1213.

11. Zhao H, et al. (2015) PARP1- and CTCF-Mediated Interactions between Active and Repressed Chromatin at the Lamina Promote Oscillating Transcription. Mol. Cell. 59(6):984–997.

12. Kim YH, et al. (2018) Rev-erbalpha dynamically modulates chromatin looping to control circadian gene transcription. Science 359(6381):1274–1277.

13. Mermet J, et al. (2018) Clock-dependent chromatin topology modulates circadian transcription and behavior. Genes. Dev. 32(5-6):347–358.

14. Shafer OT, Helfrich-Forster C, Renn SC, & Taghert PH (2006) Reevaluation of Drosophila melanogaster’s neuronal circadian pacemakers reveals new neuronal classes. J. Comp. Neurol. 498(2):180–193.

15. Dubowy C & Sehgal A (2017) Circadian Rhythms and Sleep in Drosophila melanogaster. Genetics 205(4):1373–1397.

16. Hao H, Allen DL, & Hardin PE (1997) A circadian enhancer mediates PER-dependent mRNA cycling in Drosophila melanogaster. Mol. Cell. Biol. 17(7):3687–3693.

17. Menet JS, Abruzzi KC, Desrochers J, Rodriguez J, & Rosbash M (2010) Dynamic PER repression mechanisms in the Drosophila circadian clock: from on-DNA to off-DNA. Genes. Dev. 24(4):358–367.

18. Bell-Pedersen D, et al. (2005) Circadian rhythms from multiple oscillators: lessons from diverse organisms. Nat. Rev. Genet. 6(7):544–556.

19. Yoo SH, et al. (2004) PERIOD2 :: LUCIFERASE real-time reporting of circadian dynamics reveals persistent circadian oscillations in mouse peripheral tissues. P. Natl. Acad. Sci. USA 101(15):5339–5346.

20. McDonald MJ & Rosbach M (2001) Microarray analysis and organization of circadian gene expression Drosophila. Cell 107(5):567–578.

21. Zhang R, Lahens NF, Ballance HI, Hughes ME, & Hogenesch JB (2014) A circadian gene expression atlas in mammals: Implications for biology and medicine. P. Natl. Acad. Sci. USA 111(45):16219–16224.

22. Duong HA, Robles MS, Knutti D, & Weitz CJ (2011) A molecular mechanism for circadian clock negative feedback. Science 332(6036):1436–1439.

23. Curtin KD, Huang ZJ, & Rosbash M (1995) Temporally regulated nuclear entry of the Drosophila period protein contributes to the circadian clock. Neuron 14(2):365–372.

24. Shafer OT, Rosbash M, & Truman JW (2002) Sequential nuclear accumulation of the clock proteins period and timeless in the pacemaker neurons of Drosophila melanogaster. J. Neurosci. 22(14):5946–5954.

25. Meyer P, Saez L, & Young MW (2006) PER-TIM interactions in living Drosophila cells: an interval timer for the circadian clock. Science 311(5758):226–229.

26. Shaner NC, et al. (2013) A bright monomeric green fluorescent protein derived from Branchiostoma lanceolatum. Nat. Methods 10(5):407–409.

27. Cranfill PJ, et al. (2016) Quantitative assessment of fluorescent proteins. Nat. Methods 13(7):557–562.

28. Konopka RJ & Benzer S (1971) Clock mutants of Drosophila melanogaster. P. Natl. Acad. Sci. USA 68(9):2112–2116.

29. Gummadova JO, Coutts GA, & Glossop NR (2009) Analysis of the Drosophila Clock promoter reveals heterogeneity in expression between subgroups of central oscillator cells and identifies a novel enhancer region. J. Biol. Rhythms 24(5):353–367.

30. Han C, Jan LY, & Jan YN (2011) Enhancer-driven membrane markers for analysis of nonautonomous mechanisms reveal neuron-glia interactions in Drosophila. P. Natl. Acad. Sci. USA 108(23):9673–9678.

31. Renn SCP, Park JH, Rosbash M, Hall JC, & Taghert PH (1999) A pdf neuropeptide gene mutation and ablation of PDF neurons each cause severe abnormalities of behavioral circadian rhythms in Drosophila. Cell 99(7):791–802.

32. Grima B, Chelot E, Xia RH, & Rouyer F (2004) Morning and evening peaks of activity rely on different clock neurons of the Drosophila brain. Nature 431(7010):869–873.

33. Shin Y & Brangwynne CP (2017) Liquid phase condensation in cell physiology and disease. Science 357(6357).

34. HunterEnsor M, Ousley A, & Sehgal A (1996) Regulation of the Drosophila protein timeless suggests a mechanism for resetting the circadian clock by light. Cell 84(5):677–685.

35. Chen W, Werdann M, & Zhang Y (2018) The auxin-inducible degradation system enables conditional PERIOD protein depletion in the nervous system of Drosophila melanogaster. FEBS J. 285(23):4378–4393.

36. Henry GL, Davis FP, Picard S, & Eddy SR (2012) Cell type-specific genomics of Drosophila neurons. Nucleic Acids Res. 40(19):9691–9704.

37. Price JL, et al. (1998) double-time is a novel Drosophila clock gene that regulates PERIOD protein accumulation. Cell 94(1):83–95.

38. Muskus MJ, Preuss F, Fan JY, Bjes ES, & Price JL (2007) Drosophila DBT lacking protein kinase activity produces long-period and arrhythmic circadian behavioral and molecular rhythms. Mol. Cell. Biol. 27(23):8049–8064.

39. Bindels DS, et al. (2017) mScarlet: a bright monomeric red fluorescent protein for cellular imaging. Nat. Methods 14(1):53–56.

40. Houl JH, Yu WJ, Dudek SM, & Hardin PE (2006) Drosophila CLOCK is constitutively expressed in circadian oscillator and non-oscillator cells. J. Biol. Rhythms 21(2):93–103.

41. Liu TX, Mahesh G, Houl JH, & Hardin PE (2015) Circadian Activators Are Expressed Days before They Initiate Clock Function in Late Pacemaker Neurons from Drosophila. J. Neurosci. 35(22):8662–8671.

42. Towbin BD, et al. (2012) Step-wise methylation of histone H3K9 positions heterochromatin at the nuclear periphery. Cell 150(5):934–947.

43. Yao J, Fetter RD, Hu P, Betzig E, & Tjian R (2011) Subnuclear segregation of genes and core promoter factors in myogenesis. Genes. Dev. 25(6):569–580.

44. Fung JC, Marshall WF, Dernburg A, Agard DA, & Sedat JW (1998) Homologous chromosome pairing in Drosophila melanogaster proceeds through multiple independent initiations. J. Cell Biol. 141(1):5–20.

45. Goldberg M, et al. (1998) Interactions among Drosophila nuclear envelope proteins lamin, otefin, and YA. Mol. Cell. Biol. 18(7):4315–4323.

46. Barton LJ, Soshnev AA, & Geyer PK (2015) Networking in the nucleus: a spotlight on LEM-domain proteins. Curr. Opin. Cell. Biol. 34:1–8.

47. Worman HJ, Yuan J, Blobel G, & Georgatos SD (1988) A lamin B receptor in the nuclear envelope. P. Natl. Acad. Sci. USA 85(22):8531–8534.

48. Pyrpasopoulou A, Meier J, Maison C, Simos G, & Georgatos SD (1996) The lamin B receptor (LBR) provides essential chromatin docking sites at the nuclear envelope. EMBO J. 15(24):7108–7119.

49. Ye Q & Worman HJ (1996) Interaction between an integral protein of the nuclear envelope inner membrane and human chromodomain proteins homologous to Drosophila HP1. J. Biol. Chem. 271(25):14653–14656.

50. Ye QA, Callebaut I, Pezhman A, Courvalin JC, & Worman HJ (1997) Domain-specific interactions of human HP1-type chromodomain proteins and inner nuclear membrane protein LBR. J. Biol. Chem. 272(23):14983–14989.

51. Clowney EJ, et al. (2012) Nuclear Aggregation of Olfactory Receptor Genes Governs Their Monogenic Expression. Cell 151(4):724–737.

52. Solovei I, et al. (2013) LBR and lamin A/C sequentially tether peripheral heterochromatin and inversely regulate differentiation. Cell 152(3):584–598.

53. Chen CK, et al. (2016) Xist recruits the X chromosome to the nuclear lamina to enable chromosome-wide silencing. Science 354(6311):468–472.

54. Lin ST, et al. (2014) Nuclear envelope protein MAN1 regulates clock through BMAL1. Elife 3:e02981.

55. Partch CL, Clarkson MW, Ozgur S, Lee AL, & Sancar A (2005) Role of structural plasticity in signal transduction by the cryptochrome blue-light photoreceptor. Biochemistry 44(10):3795–3805.

56. Hurley JM, Larrondo LF, Loros JJ, & Dunlap JC (2013) Conserved RNA Helicase FRH Acts Nonenzymatically to Support the Intrinsically Disordered Neurospora Clock Protein FRQ. Mol. Cell. 52(6):832–843.

57. Xu HY, et al. (2015) Cryptochrome 1 regulates the circadian clock through dynamic interactions with the BMAL1 C terminus. Nat. Struct. Mol. Biol. 22(6):476–U470.

58. Cheutin T & Cavalli G (2014) Polycomb silencing: from linear chromatin domains to 3D chromosome folding. Curr. Opin. Genet. Dev. 25:30–37.

59. Wani AH, et al. (2016) Chromatin topology is coupled to Polycomb group protein subnuclear organization. *Nat*. Communications 7.

60. Rego A, Sinclair PB, Tao W, Kireev I, & Belmont AS (2008) The facultative heterochromatin of the inactive X chromosome has a distinctive condensed ultrastructure. J. Cell. Sci. 121(7):1119–1127.

61. Kosak ST, et al. (2002) Subnuclear compartmentalization of immunoglobulin loci during lymphocyte development. Science 296(5565):158–162.

62. Kohwi M, Lupton JR, Lai SL, Miller MR, & Doe CQ (2013) Developmentally Regulated Subnuclear Genome Reorganization Restricts Neural Progenitor Competence in Drosophila. Cell 152(1-2):97–108.

63. Riabinina O, et al. (2015) Improved and expanded Q-system reagents for genetic manipulations. Nat. Methods 12(3):219–223.

64. Romero P, et al. (2001) Sequence complexity of disordered protein. Proteins 42(1):38–48.

65. Cyran SA, et al. (2005) The double-time protein kinase regulates the subcellular localization of the Drosophila clock protein period. J. Neurosci. 25(22):5430–5437.

66. So WV & Rosbash M (1997) Post-transcriptional regulation contributes to Drosophila clock gene mRNA cycling. EMBO J. 16(23):7146–7155.

